# A photoswitchable ligand targeting β_1_-adrenoceptor enables light-control of the cardiac rhythm

**DOI:** 10.1101/2022.03.06.483174

**Authors:** Anna Duran-Corbera, Melissa Faria, Yuanyuan Ma, Eva Prats, André Dias, Juanlo Catena, Karen L. Martinez, Demetrio Raldua, Amadeu Llebaria, Xavier Rovira

## Abstract

Catecholamine-triggered β-adrenoceptor (β-AR) signaling is essential for the correct functioning of the heart. Although both β_1_- and β_2_-AR subtypes are expressed in cardiomyocytes, drugs selectively targeting β_1_-AR have proven this receptor as the main target for the therapeutic effects of beta blockers in heart. Here, we report a new strategy for the spatiotemporal control of β_1_-AR activation by means of light-regulated drugs with a high level of β_1_-/β_2_-AR selectivity. All reported molecules allow for an efficient real time optical control of receptor function in vitro. Moreover, using confocal microscopy we demonstrate that the binding of our best hit, pAzo-2, can be reversibly photocontrolled. Strikingly, pAzo-2 also enables a dynamic cardiac rhythm management on alive zebrafish larvae using light, thus highlighting the therapeutic and research potential of the developed photoswitches. Overall, this work provides the first proof of precise control of the therapeutic target β_1_-AR in native environments using light.

## Introduction

Beta-adrenoceptors (β-AR) are class A G protein-coupled receptors (GPCRs) endogenously activated by the catecholamines adrenaline or noradrenaline, which regulate a variety of biological functions.^[1–3]^ Their crucial role in the regulation of cardiac function and the respiratory system, among others, has signaled them as classical pharmacological targets.^[4]^ β-AR are divided in three subtypes, β_1_-, β_2_- and β_3_-AR. All β-AR mainly induce the production of cAMP from ATP through Gs coupling.^[1,3]^ However, their biological effects are noticeably different, majorly due to their different localization in the body. β_1_-adrenoceptors, which regulate the cardiac output, are mainly located in the heart and cerebral cortex.^[5,6]^ In contrast, β_2_-AR are prevalent in the respiratory system and cerebellum and control smooth muscle relaxation processes.^[1,7]^ Finally, β_3_-AR are located in the adipose tissue and control metabolic processes such as the regulation of lipolysis and thermogenesis.^[8]^ Even though the presence of each receptor subtype is predominant in specific organs, their expression can be found in other tissues. Moreover, co-expression of different subtypes is a common phenomenon.^[9]^ Indeed, both β_1_- and β_2_-AR are expressed in cardiomyocytes and work synergistically to regulate myocardial contractility.^[5]^ Nonetheless, several studies have described β_1_-AR as the receptor with therapeutic significance in cardiac diseases compared to β_2_-AR.^[10,11]^However, the poor selectivity of beta-blockers, especially between cardiac β_1_-AR and respiratory β_2_-AR, constitutes a problem for the treatment of patients with both cardiac and respiratory conditions.^[12]^ Therefore, β_1_/β_2_-selectivity in beta-blockers is critical and an appropriate selection of the drug is sometimes challenging for the correct management of patients with respiratory and cardiac pathologies, which would benefit from more selective treatments targeting specific receptors in restricted locations of the body.

GPCR pharmacology has classically been tackled from a mono-dimensional approach, which might be inefficient considering the complexity of the signaling pathways that can be involved in a particular disease. To overcome this limitation, research has put the focus on the development of novel molecular tools that allow dynamic control of GPCR activity. Light constitutes a powerful tool to study biological systems, as it can be applied with unparalleled spatial and temporal precision. In this context, the development of molecules and techniques regulated with light has increasingly gained popularity in the study of GPCRs. One of these approaches, named photopharmacology, involves the development of small molecules with photochromic properties. ^[13–15]^ Light application is expected to trigger a change in the geometry, polarity, and electron density of the light-sensitive ligand, which is expected to alter its pharmacological properties.^[13,16,17]^ In consequence, the use of photochromic ligands allows modulating the activation of a targeted receptor with spatiotemporal precision. This innovative technique has been widely applied for research purposes, and several photochromic ligands have been described in the literature allowing the optical control of a wide variety of GPCRs.^[18–21]^Moreover, interesting pharmacological properties have emerged from the use of photochromic molecules, such as ligands with opposing functional properties (agonists/antagonists) depending on the applied light, among others.^[22,23]^ These studies have provided valuable information on the mechanisms of receptor function and activation, the importance of receptor localization for its function, and the relevance of the temporal dimension, among others.^[24]^ Moreover, some of the developed ligands have also been used for the optical control of physiological functions in wild-type living animals, demonstrating the potential of photopharmacology for physiological studies in native environments, including a recent example in cardiac photopharmacolgy.^[19,25]^ Although the GPCR photopharmacological toolbox contains a variety of pharmacological and chemical strategies, only a few studies have been focused on β-AR^[26–28]^ and, surprisingly, there are no light-regulated ligands for the modulation β_1_-AR reported in the literature so far.

In this study, we present a new strategy for the development of photoswitchable ligands targeting β_1_-AR. The design of these ligands is based on cardioselective known betablockers, which are characterized by a para-aromatic substitution pattern.^[29]^ Introducing the oxyaminoalcohol moiety in *para*-with respect to the azo bridge yielded light-regulated ligands with a high degree of β_1_-/β_2_-AR selectivity. The azobenzene molecules demonstrate a temporal control of the receptor activity in cell cultures. Moreover, these photopharmacological tools are compatible with confocal microscopy methods that are used herein to show evidence of receptor binding reversibility. Strikingly, we demonstrate that important physiological processes such as the heart rhythm can be dynamically controlled with light in alive zebrafish larvae using these photopharmacological agents.

## Results and Discussion

### Design and synthesis

It is well described that β_1_-AR and β_2_-AR are highly similar in sequence and structure.^[30]^ However, despite their high homology, numerous agonists, and antagonists selective for β_1_-AR or β_2_-AR have been discovered.^[30–32]^ An evaluation of the chemical structures of β_1_-AR selective antagonists highlighted the presence of an ethanolamine backbone linked to an aromatic unit through an oxymethylene bridge. The ethanolamine moiety is essential to obtain molecules with β-AR activity, as it forms interactions with key residues in the orthosteric pocket of the target receptor. On the other hand, the oxymethylene linker is a recurrent molecular particularity on the structure of β-AR antagonists, although it can also be found in the structure of several partial agonists. We have recently employed this strategy to obtain nanomolar antagonists for β_2_-AR.^[28]^ Therefore, these structural determinants were conserved in the proposed molecules. To confer selectivity for β_1_-AR in front of β_2_-AR, we explored an interesting feature found in many β_1_-AR selective antagonists, where the oxyaminoalcohol substructure is repeatedly positioned in *para*-with respect to the other substituents of the phenyl ring (Figure 1). Consequently, two *para*-substituted azobenzenes (p-ABs), named **Parazolol-1** (**pAzo-1**) and **Parazolol-2** (**pAzo-2**) were designed as potential β_1_-AR selective photoswitchable ligands (Figure 1). **pAzo-1**, which presents a *para*-monosubstituted azobenzene, was proposed due to its synthetic accessibility. In **pAzo-2**, a methoxy group was introduced in the 4-position of the phenyl ring. The introduction of a p-electron-donating group (EDG) is expected to shift the π-π* transition towards the visible range of the spectrum. Moreover, **Photoazolol-3** (**PLZ-3**), a compound previously described as not active for β_2_-AR with an acetamide group in the 4-position of the phenyl ring^[28]^ is also analyzed in this study for its activity towards β_1_-AR.

**Figure 1.**
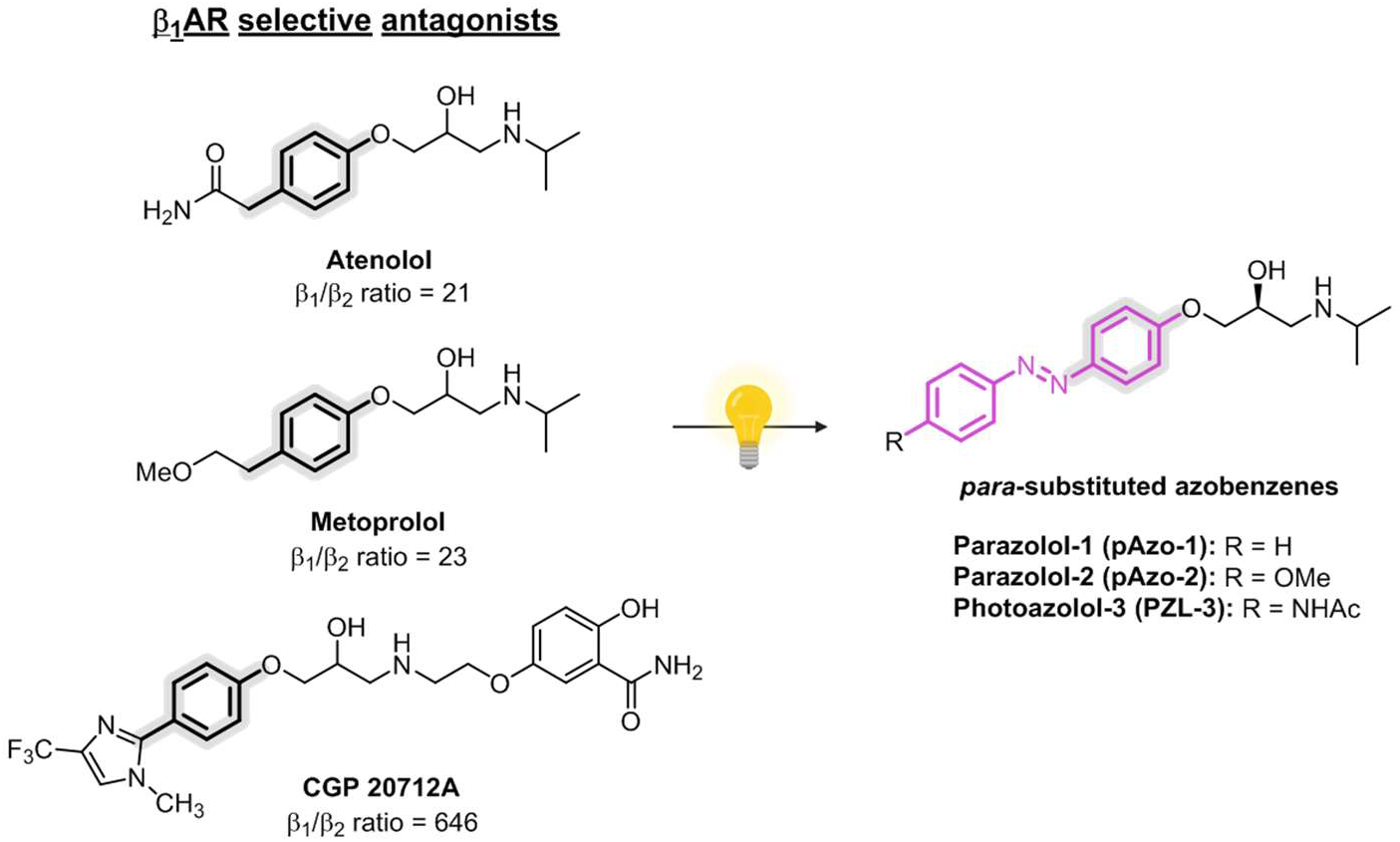
Molecular design of selective photoswitches targeting β_1_-AR. Left, prototypical β_1_-adrenoceptor selective ligands and their reported β_1_/β_2_ ratios.^[32]^ Right panel, photoswitchable ligands designed through the application of the azologization strategy.

To produce **pAzo-2**, direct diazotization of p-methoxyaniline **1** followed by reaction with phenol, which yielded intermediate **3** (Scheme 1). Unsubstituted phenol intermediate **2** was obtained commercially. Both azobenzenes (**2** and **3**) were O-alkylated by reaction with (R)-epichlorohydrin using butanone as a solvent, which proceeded with inversion of configuration to yield (S)-oxiranes **4** and **5**.^[33]^ The resulting epoxides were finally opened by nucleophilic attack of isopropylamine, and the desired products (**pAzo-1** and **pAzo-2**) were obtained.

**Scheme 1.**
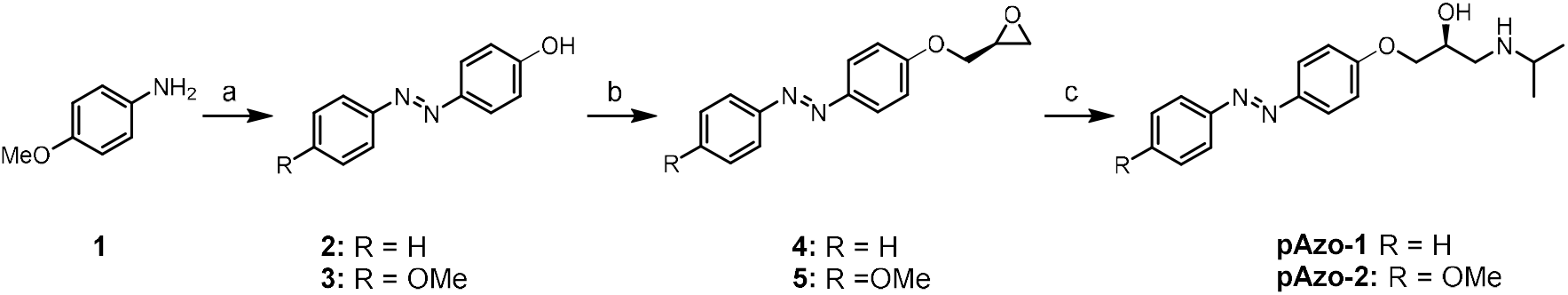
Synthesis of photoswitchable ligands selectively targeting β_1_-AR.^a^. ^a^ Reagents and conditions: (a) (I) NaNO_2_, aq HCl, 0ºC, 5 min; (II) Phenol, aq NaOH, 0ºC, 30 min 79%; (b) (R)-Epichlorohydrin, K_2_CO_3_, butanone, reflux, 24-48h, 90%; (c) i-PrNH_2_, 2-12h, r.t or MW, 24-62%.

### Photochemical characterization of azobenzenes

Following the synthesis of **Parazolols** (**pAzos**), we focused our attention on their photochemical properties. The presence of the azobenzene moiety in both ligands allows the existence of the compounds in two distinct isomeric forms, which can be reversibly interconverted through the application of light by direct excitation of their azoaromatic units (Figure 2A and S1A).

**Figure 2.**
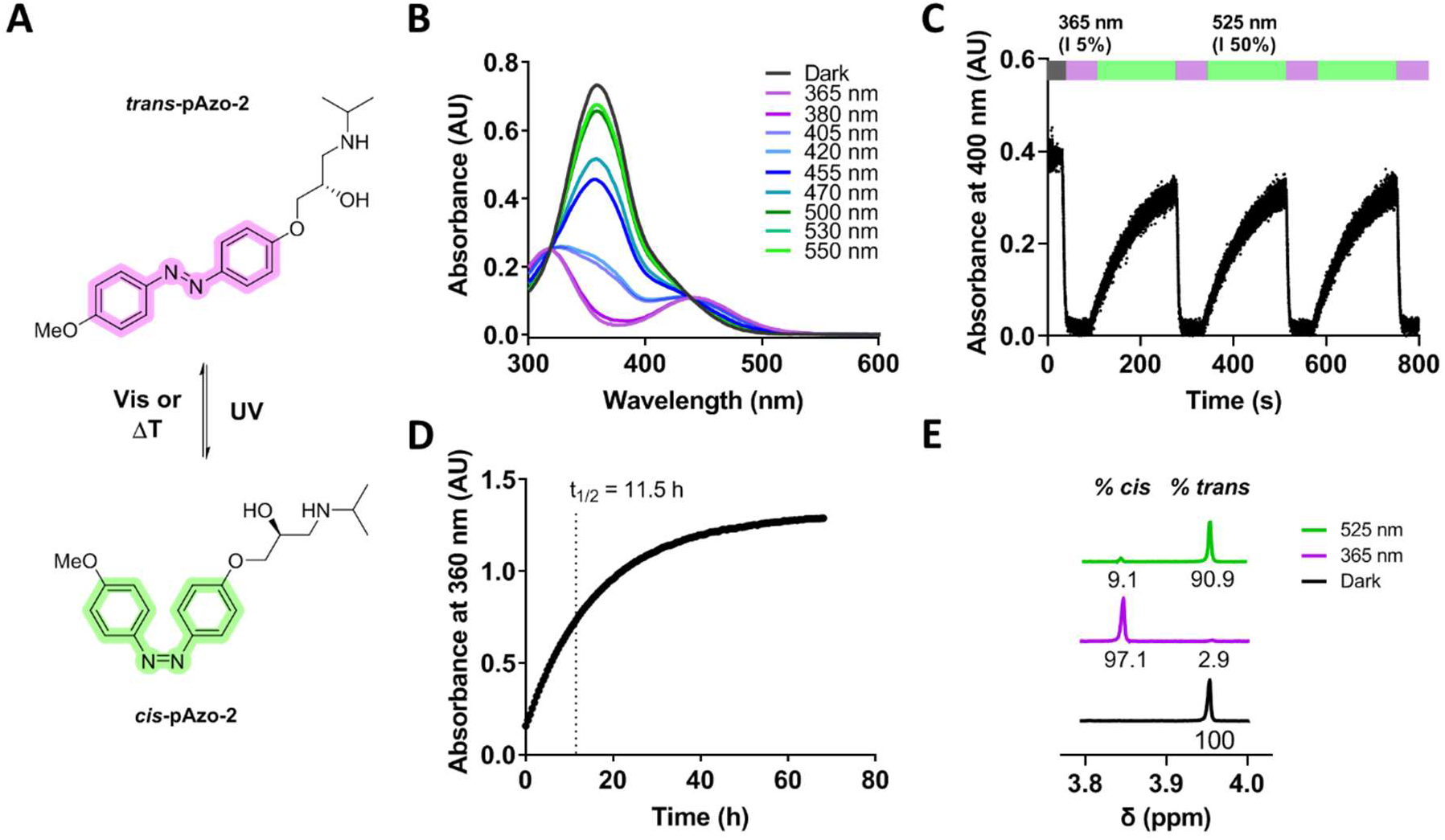
Photochemical characterization of pAzo-2. (A) 2D chemical structures of the two photoisomers of azobenzene **pAzo-2**. (B) UV-Vis absorption spectra of a 50 μM solution of **pAzo-2** in Epac buffer (0.5% DMSO) under different light conditions. (C) Multiple *cis/trans* isomerization cycles triggered by application of 365 nm (intensity set at 5%) and 525 nm (intensity set at 50%); CoolLED light system was used to apply light while absorbance at 400 nm was continuously measured. (D) Half-lifetime estimation of cis-**pAzo-2** at 25 °C in EPAC buffer (0.5% DMSO); absorbance was measured at 360 nm after 3 min of continuous illumination with 365 nm light. (E) Photostationary state (PSS) quantification by ^1^H-NMR.

To determine the optimal photoisomerization wavelengths, UV-Vis spectra of the two compounds were recorded in dark and after continuous illumination with different wavelengths for 3 minutes (Figures 2B and S1B). The absorption spectra of the *trans* isomer displayed an intense absorption band at 346 nm and 358 nm for **pAzo-1** and **pAzo-2**, respectively (Table 1). Additionally, both spectra showed a broad shoulder between 400-450 nm that corresponds to the n-π* transition, forbidden by symmetry. It is worth noting that the additional EDG in **pAzo-2** causes an energetic decrease of the π-π* transition, resulting in a bathochromic shift of the absorption band and higher overlapping with the n-π* transition (Figure 2B). Efficient excitation of the compounds to their *cis* isomers was achieved upon illumination with 365 nm for **pAzo-1** and 365/380 nm for **pAzo-2**. Back isomerization to the thermodynamically stable *trans* configuration was achieved upon illumination at 530 nm for both compounds (Figures 2B and S1B).

**Table 1.** Photochemical properties of azobenzenes pAzo-1 and pAzo-2^a^

| Compound | $\lambda_{\text{trans}}^{\text{a}}$<br>(nm) | $\lambda_{\text{cis}}^{\text{a}}$<br>(nm) | $t_{1/2}^{\text{a}}$ (h) | PSS<sub>365</sub><sup>b</sup><br>(% *cis*) | PSS<sub>530</sub><sup>b</sup><br>(% *trans*) | $T_{\text{trans} \rightarrow \text{cis}}^{\text{a,c}}$<br>(s) | $T_{\text{cis} \rightarrow \text{trans}}^{\text{a,d}}$<br>(s) |
| --- | --- | --- | --- | --- | --- | --- | --- |
| **pAzo-1** | 346 | 430 | 38.3 | 95.2 | 76.3 | 7.4 | 122.1 |
| **pAzo-2** | 358 | 440 | 11.5 | 97.1 | 90.9 | 2.9 | 69.0 |

<sup>a</sup> Determined at 50 μM in aqueous buffer + 0.5 % DMSO, 25 °C. <sup>b</sup> PSS state areas were determined at 12 °C by <sup>1</sup>H-NMR after illumination (365/525 nm) of a 100 μM sample in D<sub>2</sub>O. <sup>c</sup> 365 nm light was applied to trigger excitation to the *cis* isomer; The CoolLED light system set at 5% intensity was used. <sup>d</sup> 525 nm light was applied to trigger back-isomerization using the CoolLED set at 50% intensity.

Furthermore, both compounds showed reversible photoisomerization through the application of several light cycles continuously recorded (365/525 nm), which allowed the determination of isomerization rates (⊤) and confirmed the stability of the two compounds to prolonged illuminations (Figure 2C, S1C, and Table 1). Thermal relaxation of the *cis* isomers in an aqueous solution was also evaluated at 25°C (Figure 2D and S1D). Both compounds showed long cis-state thermal stability, with half-life times of 38.3 h and 11.5 h for **pAzo-1** and **pAzo-2**, respectively. Compound **pAzo-2**, which contains an additional methoxy group in the 4-position of the phenyl ring, showed a noticeably faster thermal relaxation compared to the monosubstituted *p*-AB **pAzo-1**.^[34]^

Finally, relative concentrations of the two isomers in equilibrium after illumination with 365 nm and 525 nm (PSS_365_ and PSS_525_) were determined by ^1^H NMR spectroscopy (Figure 2E, S1E, and Table 1). Upon illumination at 365 nm for 3 minutes, all signals present in the dark spectra were shifted, which confirmed the excitation of the compounds to the *cis* isomer. Both compounds showed almost quantitative conversion (< 95%) to the *cis* isomer upon illumination at 365 nm. Illumination at 525 nm triggered the *cis→trans* isomerization in both compounds. For **pAzo-2**, 90.9% of the thermostable *trans* isomer was recovered upon illumination. Nevertheless, the photoinduced back-isomerization of *cis*-**pAzo-1** was not as efficient, with only 76.3% of the *trans* isomer quantified at PSS_525_. This lower efficiency appears as a consequence of the n-π* band overlapping found between the absorbance spectra of the two photoisomers of **pAzo-1** (Figure S1B).

### pAzo compounds reversibly modulate β_1_-AR activity as partial agonists

Following the photochemical characterization of **pAzo-1** and **pAzo-2**, light-dependent pharmacological properties of the two photochromic ligands towards β_1_- and β_2_-AR were evaluated in cultured cells. In the activity tests we also included **Photoazolol-3** (**PZL-3**), a recently reported adrenergic photochromic ligand with a *p*-acetamido substituent.^[28]^ This ligand, which also displays the p-AB scaffold proposed to selectively target β_1_-AR (Figure 1), was found to have negligible inhibitory potency against β_2_-AR. Nevertheless, the pharmacological activity of its two photoisomers was never assessed against β_1_-AR, thus raising interest for its potential as a β_1_-AR selective photoswitchable ligand. ^[28]^

β_1_-AR photopharmacological properties of p-ABs were evaluated in HEK293 iSNAP β_1_AR H188 cells, which express β_1_-AR upon induction with doxycycline. A double stable cell line was generated with an EPAC S^H188^ CFP-YFP FRET biosensor,^[35]^ which allowed to continuously monitor intracellular cAMP levels. Considering that this cell system allows to control β_1_-AR expression levels, we optimized functional assays after 24h induction (Figure S2). The lower level of expression for which the reference agonist cimaterol remained equally potent was taken. That was achieved by adding 0.01 μM of the induction agent Doxycycline.

Results showed that *trans* isomers of all p-ABs displayed good agonistic activity with a β_1_-AR EC50 in the nanomolar range (Figure 3A-C). Application of violet light (380 nm) triggered a significant decrease in the EC50 of the photoswitchable ligands, which appointed them as *trans-on* compounds, that is, the *trans* isomer is more active than the *cis* isomer (Figure 3A-C and Table 2). Moreover, the tested photoswitches displayed very good light-dependent modulation of β_1_-AR, with photoinduced potency shifts (PPS) ranging from 20- to 83-fold (Table 1). Among the three ligands, **pAzo-2** presented a particularly promising behavior, as it allowed modulation of the activation state of β_1_-AR from 100% to almost 0% through the application of light at concentrations around 100 nM. On the other hand, **PZL-3** displayed lower agonistic potency compared to **pAzos**, with an approximate 10-fold decrease in the EC_50_ values (Table 2).

**Figure 3.**
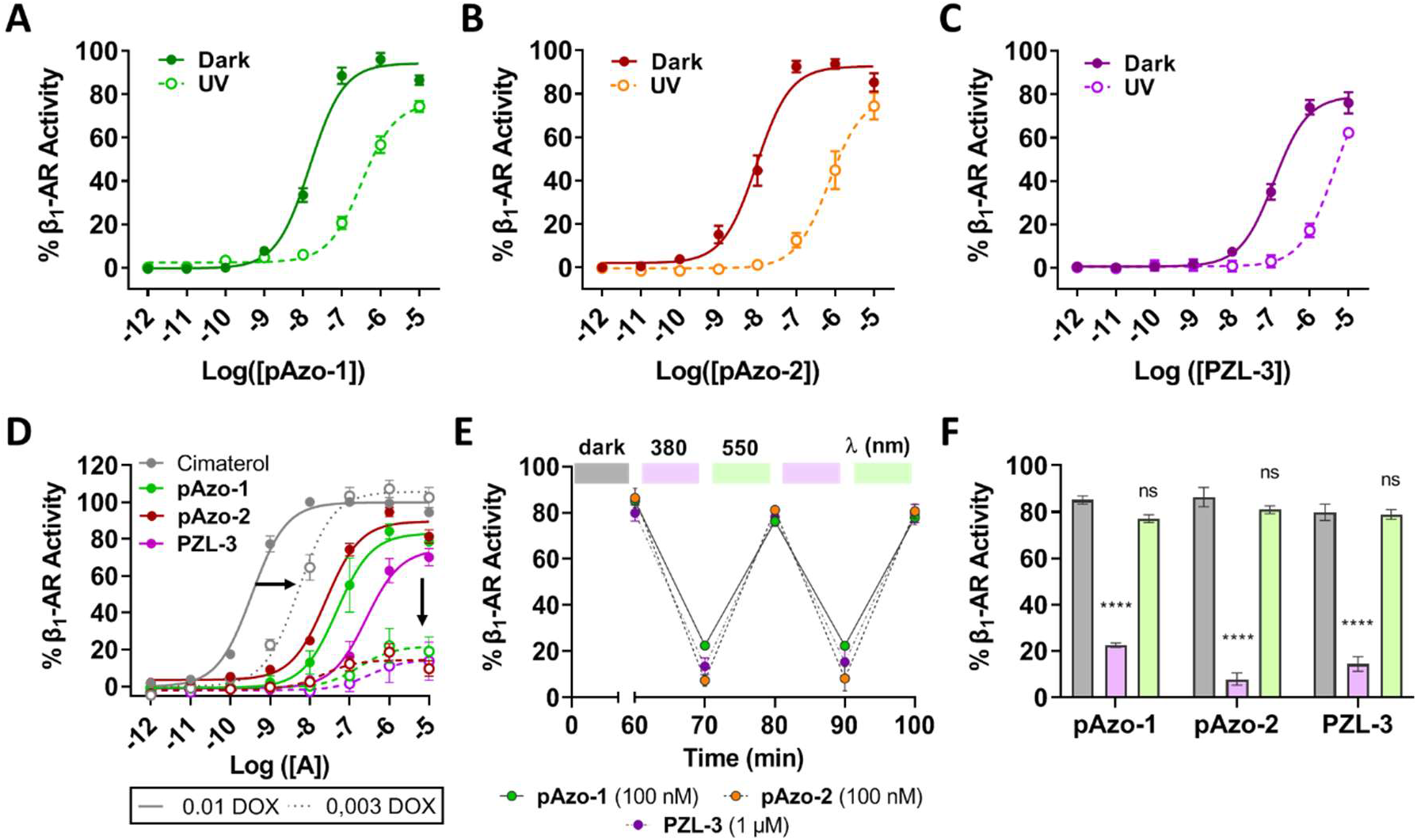
Light-dependent modulation of β_1_-AR functional response. Dose-response curves of **pAzo-1** (A), **pAzo-2** (B), and **PZL-3** (C) in the dark (solid line) and after the application of constant violet light at 380 nm (dotted line). (D) Dose-response curves of cimaterol (in grey), *trans*-**pAzo-1** (green), *trans*-**pAzo-2** (dark red), and *trans*-**PZL-3** (purple) performed in HEK293 iSNAP β_1_AR H188 cells with higher and lower levels of receptor expression (induced by 0.01μM and 0.003 μM Doxycycline (DOX)). (E) Time-course quantification of intracellular cAMP in the presence of **pAzo-1** (green dots), **pAzo-2** (orange dots), and **PZL-3** (purple dots). Purple and green boxes correspond to 10 min illumination breaks using 380 and 550 nm lights, respectively. (F) Receptor activity values measured for the different light conditions. Data are shown as the mean ± SEM of three to five independent experiments performed in duplicate. Statistical differences are denoted for adjusted p-values as follows: ****p<0.0001.

**Table 2.** Pharmacological data of pAzo-1 and pAzo-2 towards β_1_-AR.

| Cmpd | DARK |  |  | LIGHT |  |  |  |  |
| --- | --- | --- | --- | --- | --- | --- | --- | --- |
| | EC<sub>50</sub> (nM) | SEM | $\beta_1/\beta_2$ ratio<sup>a</sup> | EC<sub>50</sub> (nM) | SEM | $\beta_1/\beta_2$ ratio<sup>a</sup> | PPS<sup>b</sup> | SEM |
| **pAzo-1** | 15.14**** | 1.42 | 161.8 | 331.13 | 77.76 | 8.6 | 20.08 | 6.06 |
| **pAzo-2** | 8.79**** | 2.89 | 187.7 | 762.08 | 169.61 | 2.3 | 83.03 | 39.06 |
| **PZL-3** | 120.50**** | 12.46 | 93.8 | 4581.41 | 1026.81 | 3.1 | 37.67 | 9.55 |

<sup>a</sup> A ratio of 1 implies no selectivity towards $\beta_1$ -AR. <sup>b</sup> PPS refers to Photoinduced Potency Shift, which is the relation between the measured EC<sub>50</sub> in light and dark conditions respectively. Statistical differences from light EC<sub>50</sub> values are denoted for adjusted p values as follows: \*\*\*\*p<0.0001.

To evaluate if the developed azobenzenes are full or partial agonists, dose-response curves of cimaterol and the three photoswitchable ligands (**pAzos** and **PZL-3**) were performed in HEK293 iSNAP β_1_AR H188 cells with two different β_1_-AR expression levels (induced with 0.01 μM and 0.003 μM doxycycline for 24h, respectively) (Figure 3D).^[36]^Results showed that the use of cells with lower expression levels only shifted the doseresponse curve of full agonist cimaterol, thus significantly increasing its measured EC_50_ in this system (Figure 3D). Meanwhile, changes in the expression levels of β_1_-AR did not alter the potency of the tested compounds (EC_50_ values were unaltered, Table S2) but significantly affected their efficacy. Indeed, E_max_ values were greatly reduced when doseresponse curves of **pAzos** and **PZL-3** were registered in cells with lower expression of β_1_-AR (Table S2). To further confirm that the observed effects are related to modulation of β_1_-AR, dose-response curves of the tested photoswitches with a constant concentration of a β_2_-AR selective antagonist (ICI 118511) were performed with the two different β_1_-AR expression levels. No significant changes were observed between the curves obtained for the three ligands in the presence or absence of the selective antagonist (Figure S3). Overall, these results evidence that the three *p*-AB behave as partial agonists with low efficacy towards β_1_-AR at low levels of expression, similar to those we expect to find in native systems.^[36]^

On the other hand, the functional β_2_-AR activity of the two **pAzos** was assessed using HEK293 H188 M1 cells, following a protocol recently reported in the literature.^[28]^ Briefly, inhibitory dose-response curves of the ligands were obtained with a constant concentration of cimaterol (3 nM), both in the dark and upon illumination at 380 nm. Both **pAzo-1** and **pAzo-2** displayed low inhibitory potency against β_2_-AR (IC_50_ values within the μM range), and no significant light-induced changes in their pharmacological behavior (Figure S4 and Table S1). These results were in good agreement with the pharmacological properties reported for **PZL-3** towards β_2_-AR.^[28]^

Therefore, the three p-ABs displayed β_1_/β_2_ selectivity ratios ranging from 93.8 to 161.8 (Table 2), at the level of β_1_-AR selective ligands atenolol and metoprolol, which are approved drugs considered cardioselective (Figure 1).^[31]^ On the other hand, the *cis* isomers of the three compounds showed moderate modulation of the two adrenoceptor subtypes and displayed lower β_1_/β_2_ selectivity ratios, ranging from 2.3 to 8.6 (Table 2). The fact that good selectivity is achieved when the compounds were disposed in their longer isomeric form was in good agreement with the recently reported molecular mechanisms governing β_1_/β_2_ subtype selectivity.^[32,37,38]^ Both β_1_-AR and β_2_-AR present the same residues at the orthosteric site, only some residues at the edge of the pocket are different. Thus, long ligands may directly interact with these residues resulting in differential modulation of the two receptor subtypes.^[39]^

Finally, studies were performed to assess the dynamic and reversible modulation of the target receptor with light in living cells. In these experiments, the tested compounds enabled good modulation of the activation state of β_1_-AR *in vitro* through the application of light cycles (Figure 3E and F). Results showed that the three *p*-ABs (**pAzos** and **PZL-3**) displayed similar agonistic effects in the dark (80-90% receptor activation). When violet light was applied to cells, a significant decrease in the activation state of β_1_-AR was measured for the three azobenzenes, implying that their *cis* isomers display reduced agonism, which was consistent with their *trans-on* nature. Back-isomerization to their *trans* state was efficiently achieved through the application of green light, which restored initial activation levels of β_1_-AR. Therefore, the developed ligands show good and reversible photopharmacological properties against β_1_-AR *in vitro* (Figure 3E and F).

To sum up, we have successfully developed the first photoswitchable ligands that enable a reversible modulation of β_1_-AR through the application of light. The three p-ABs are *trans*-on compounds with partial agonist pharmacology and potencies in the nanomolar range. The application of violet light (380 nm) triggered excitation of the ligands to their *cis* isomeric forms, which displayed significantly lower agonistic activities towards β_1_-AR. In particular, the *p* 4-methoxy-substituted compound (**pAzo-2**) was identified as the best hit, as it showed the highest agonistic potency and photoinduced shift, as well as better β_1_-/β_2_-AR selectivity. Moreover, it allowed to dynamically modulate the activation state of β_1_-AR from 90% in its *trans* state to approximately 6% upon light application (Figure 3E and F). In consequence, **pAzo-2** is a particularly interesting photoswitchable ligand, which enables an almost complete ON/OFF modulation of β_1_-AR with the external application of different light periods.

### pAzo-2 binding photocontrol can be precisely monitored using confocal microscopy

Optical imaging is a powerful technique to interrogate biological systems, validate pharmacological approaches, and study molecular processes. With the twofold aim of evaluating the compatibility of the developed compounds with imaging methodologies and assessing the dynamic binding and unbinding processes of **pAzo-2** under different light conditions, a series of experiments were performed using single-cell analysis from confocal microscopy images. The method was validated to ensure the absence of fluorophore bleaching induced by the application of light to the cellular system. Control experiments demonstrated negligible influence of light application over the emitted fluorescence and confirmed the protocol compatibility with the fluorophores tagging receptors and ligands (Figure S5). Specific detection of β_1_-AR was achieved by selectively labelling the target receptors with SNAP-surface-647. The binding of **pAzo-2** to β_1_-AR preincubated with the fluorescent ligand alprenolol-green (20 nM) was measured under different light conditions. After the addition of **pAzo-2** (100 nM), the fluorescent ligand was displaced in dark conditions (Figure 4A). In contrast, after 10 min illumination with violet light the competition between the ligands for the β_1_-AR binding site completely disappeared, which was consistent with results obtained in functional assays (Figure 3E). Interestingly, binding of **pAzo-2** was recovered upon subsequent illumination of the sample with laser light at 559 nm, thus demonstrating its reversible behavior, in consistency with the described pharmacological results (Figure 2C and 3E). In addition, the amount of bound ligand in dark and light conditions was measured for a range of **pAzo-2** concentrations. The affinity of **pAzo-2** was found to be significantly influenced by the application of violet light (Figure 4D), with a 178-fold change between potencies in the dark (IC_50_ dark = 7.24nM) and after illumination (IC_50_ violet = 1.29 μM). On the other hand, the binding properties of non-photoswitchable ligand propranolol were not affected by light application, as no significant differences were detected on its affinity before and after illumination (8.67 ± 0.07 vs. 8.46 ± 0.14, respectively; Figure 4C and S6). Consequently, these results suggest that the functional differences detected between the two isomeric forms of **pAzo-2** are due to a significant reduction in the affinity of *cis*-**pAzo-2**. Moreover, the obtained results demonstrate the usefulness and compatibility of the photoswitchable ligand **pAzo-2** with common microscopy methods used in biological research. This opens the door to the development of future projects where the dynamics of β_1_-AR can be precisely explored.

**Figure 4.**
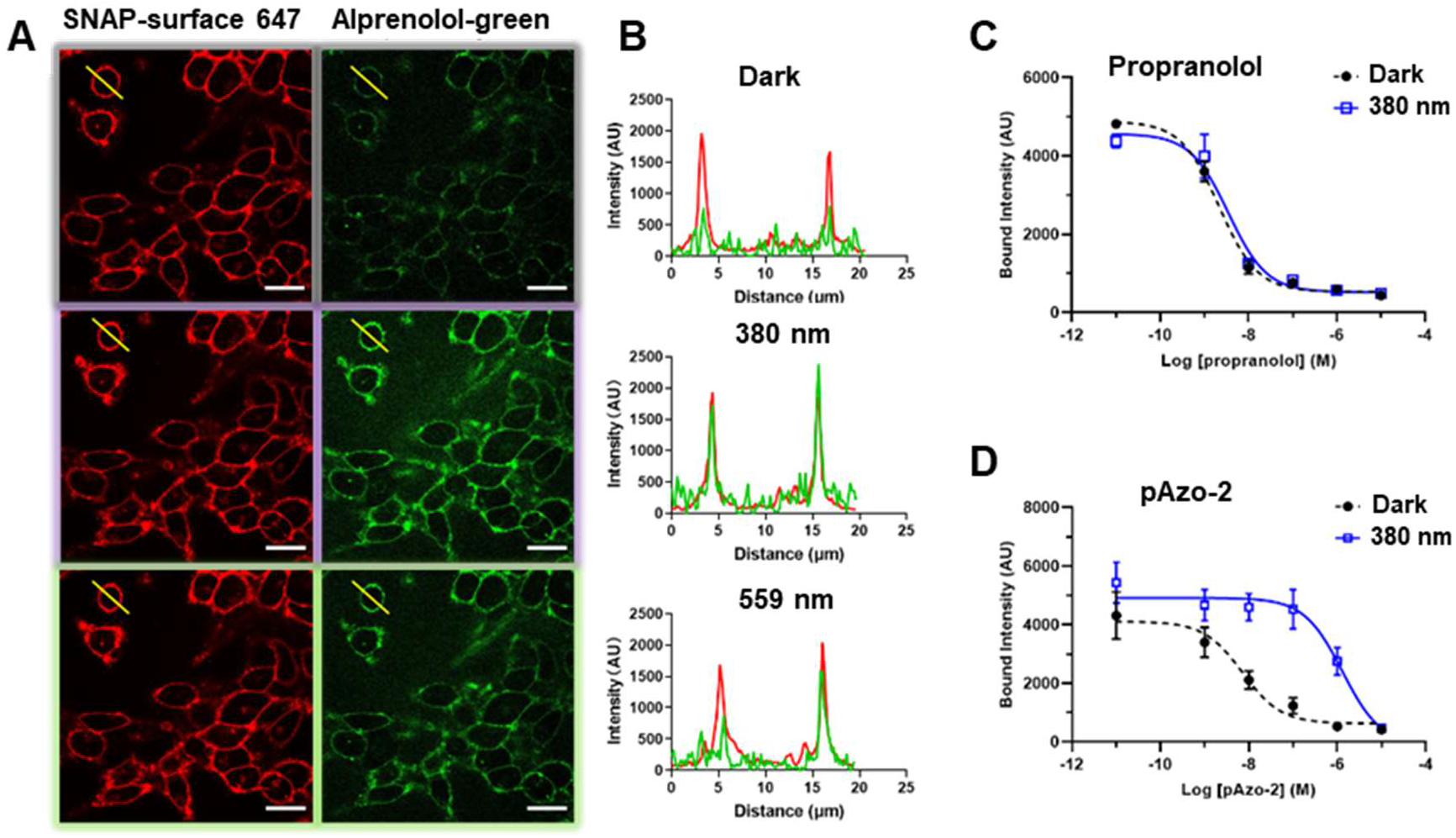
Light-dependent control of pAzo-2 binding on β_1_-AR. (A) Representative confocal fluorescence images of cells expressing SNAP-β_1_-AR labelled with SNAP-surface-647 and preincubated with alprenolol-green (20 nM) for 30 min. Cells were treated with 100 nM **pAzo-2** in dark for 1 h. Images were obtained in the dark (Top panel), after 10 min violet light exposure (Middle panel) and after 5 min green laser exposure (bottom panel). Scale bars are 20 μm. (B) Fluorescence signals measured along the yellow line in Fig. A. The receptor at the plasma membrane is shown in red and the bound alprenolol-green is shown in green for the three conditions: dark (top), violet illumination (middle) and green illumination (bottom). Competitive binding curves were extracted from confocal fluorescence images of single cells for several concentrations of propranolol (C) and **pAzo-2** (D). Compounds were added and measured in the dark (black line) and after a violet light treatment (blue line). All data are the average of three independent experiments ± SEM. The number of cells analyzed for **pAzo-2** and propranolol was 2156 and 1862, respectively.

### pAzo-2 enables a reversible light-control of the cardiac rhythm in zebrafish larvae

In order to assess the potential of the developed p-ABs for research and therapeutic applications, the capability of **pAzo-2** to modulate the cardiac rhythm through the application of light was evaluated. Zebrafish larvae (7 days post-fertilization) were exposed to different treatments (Control, 25 μM **pAzo-2** and 10 μM carvedilol) under dark conditions (1.5-2 h exposures) and the cardiac rhythm of the larvae was monitored using a microscope equipped with a camera (Figure 5A). Additionally, light-triggered effects on the cardiac rhythm were assessed. To this aim, the different experimental groups were kept in dark conditions for 1-1.5 h and thereafter illuminated for 1 min with 380 nm light. To evaluate reversibility of the drug action, a subsequent illumination of animals with 550 nm light for 1 min was performed (Figure 5B). The treatment of the larvae with trans-**pAzo-2** in the dark caused a significant reduction of the cardiac rhythm. These results are in good agreement with the cellular data obtained *in vitro*, where **pAzo-2** appears as a partial agonist with low efficacy at lower levels of expression (Figure 3D). Interestingly, when animals exposed to **pAzo-2** were illuminated with violet light at 380 nm, the measured heart rate was restored to control levels. Subsequent illumination with green light at 550 nm produced a general decrease of the heart rate in all experimental groups (Figure S7). Nonetheless, this light also reactivated **pAzo-2**, further decreasing the heart rate in comparison to the control and equivalent to the initial dark conditions (Figure 5B). Importantly, larvae treated with the non-photoswitchable antagonist carvedilol experienced a significant reduction in the cardiac rhythm, and equivalent changes were detected between the groups kept in dark and both illumination conditions (Figure 5B). These data confirm that **pAzo-2** exerts different and reversible light-triggered pharmacological effects in zebrafish larvae. Therefore, these experiments highlight the potential of the developed ABs, which enable a reversible modulation of β_1_-AR function and allow cardiac control, both through the application of ligand and light in living animals.

**Figure 5.**
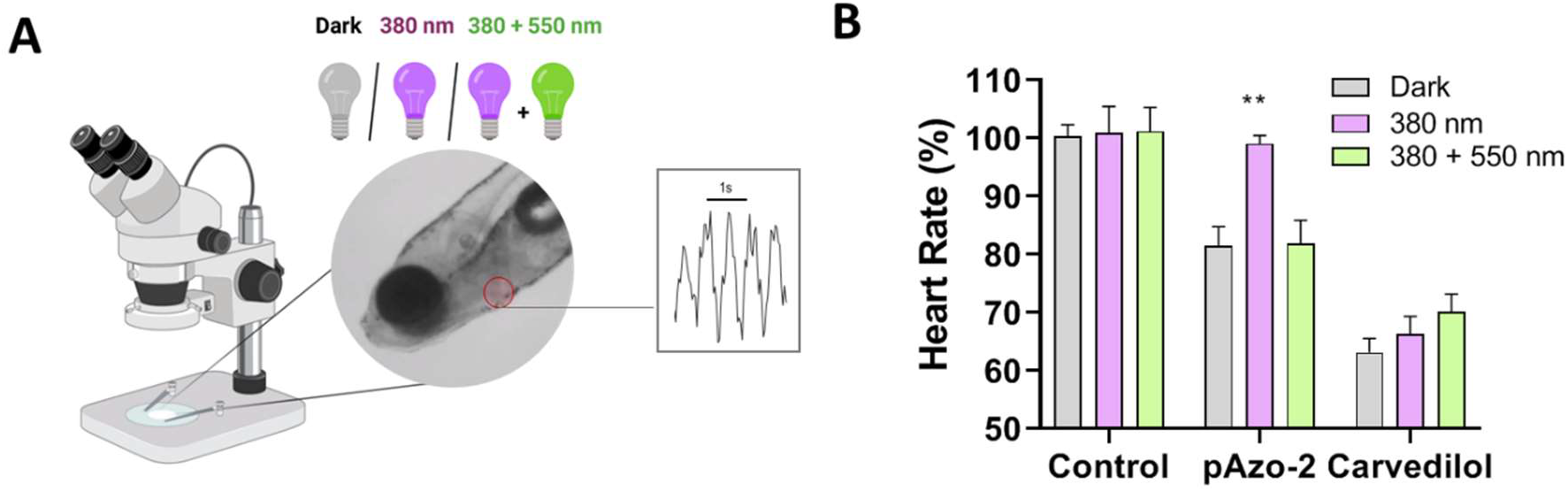
Optical modulation of the cardiac frequency by pAzo-2. (A) Protocol followed to assess cardiac modulation of through the application of **pAzo-2** light. (B) Normalized cardiac frequency of the experimental groups (control with 1% DMSO N=9-18, 25 μM **pAzo-2** N=8-14 and 10 μM carvedilol N=14-15) in the dark, after illumination with 380 nm light for 1 min and after a subsequent illumination with 550 nm light. Data are shown as the mean ± SEM of two independent experiments. Statistical differences between the different illumination conditions are denoted for adjusted p values as follows: **p<0.01

## Conclusions

In summary, we report here **pAzo-1** and **pAzo-2**, the first photoswitchable ligands targeting β_1_-AR. These molecules, which are partial agonists with nanomolar potencies in the dark, displayed high β_1_-/β_2_-AR selectivity ratios, comparable to those of marketed drugs. Upon illumination with violet light, the tested ligands showed significantly lower potencies, which could be reversed through the application of green light. These light-induced changes detected in the functional response of β_1_-AR can be attributed to a significant affinity loss for the compounds in their *cis* state, as *cis*-**pAzo-2** displayed 178-fold lower affinity values than *trans*-**pAzo-2**. Notably, we have established a β_1_-azobenzene ligand design with easy synthesis and handling. Moreover, we have developed original pharmacological cell assays to study β_1_-AR photopharmacology.

To show some of the future promising features of the best hit among the tested p-ABs, we have effectively demonstrated light-induced β_1_-AR receptor photoswitching of **pAzo-2** in confocal microscopy experiments. Using this method, real-time receptor regulation can be achieved, thus demonstrating the compatibility of this ligand with a fluorescence readout and opening new avenues to mechanistic studies. On another aspect, we have provided a proof of concept for the potential of **pAzo-2** *in-vivo*. Exposing zebrafish larvae to **pAzo-2** enabled reversible control of the cardiac rhythm through the application of light. These results account for the potential of **pAzo-2** in the study and control of cardiac physiology, where β_1_-AR plays a fundamental role. New drugs with improved performances are expected for the treatment of cardiac diseases.

Challenges for the ongoing β_1_-AR photoregulation with azobenzenes in future projects of our group will include: (i) improving ligand performances, such as red-shifting photoisomerization, (ii) understanding the molecular details of ligand-receptor interactions, (iii) performing receptor mechanistic studies and (iv) exploring therapeutic photopharmacology for β_1_-AR related diseases. These objectives can require tailoring new molecules for specific uses. Nevertheless, **pAzo-2** and related compounds are potent photoswitches with robust performance that can easily find uses to understand GPCR dynamics and cell communication, how different pathways give rise to physiological effects, and how to promote precise interaction of drugs with biological systems. Another aspect is related to in vivo applications including rodents, which will define the potential for future development of advanced drugs for cardiac diseases that can be applied systemically and locally activated in the heart for defined periods of time with light pulses.

Overall, the present work provides innovative light-regulated molecules that can be used to better understand the function and dynamics of β_1_-AR in their natural environment with high spatiotemporal precision using light.

## Experimental section

Detailed experimental procedures, synthesis methods, characterization data and original spectra are presented in Supporting Information.

## Acknowledgments

We thank Ignacio Pérez (IQAC-CSIC, Barcelona), Yolanda Pérez (IQAC-CSIC, Barcelona), Lourdes Muñoz (SimChem, IQAC-CSIC, Barcelona) and Carme Serra (SimChem, IQAC-CSIC, Barcelona) for technical support. We thank Dr. Kees Jalink (The Netherlands Cancer Institute, Amsterdam, the Netherlands) for providing the plasmids encoding for the Epac-SH188 biosensor. We thank the University of Vic-Central University of Catalonia (UVic-UCC) and Dr. Marta Otero for the material assignment which helped in some biological assays. We thank Nikos Hatzakis for access to the Olympus IX81 confocal microscope (UCPH, DK). This work was supported by Ministerio de Ciencia e Innovatión, Agencia Estatal de Investigación and ERDF-FEDER European Fund (projects CTQ2017-89222-R and PID2020-120499RB-I00) and by the Catalan government (2017 SGR 1604 and 2017-SGR-1807) to AL. XR research was financed by the Spanish Ministry of Economy, Industry and Competitiveness (SAF2015-74132-JIN). DR research was supported by “Agencia Estatal de Investigación” from the Spanish Ministry of Science and Innovation (project PID2020-113371RB-C21) and IDAEA-CSIC, Severo Ochoa Centre of Excellence (CEX2018-000794-S), which financed M.F. ADC received the support of a fellowship from “la Caixa” Foundation (ID 100010434) under the fellowship code LCF/BQ/DE18/11670012. KLM, AD and YM were supported by the Novo Nordisk Foundation (NNF20OC0064565).

## Entry for the Table of Contents

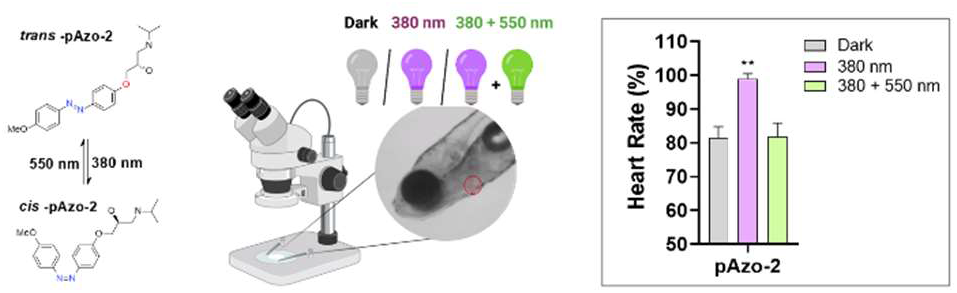
We present new photoswitchable molecules acting as partial agonists for the therapeutic β_1_-adrenoceptor (β_1_-AR). The most promising candidate, named **pAzo-2**, has a potency and β_1_/β_2_ selectivity similar to approved beta blockers. Moreover, **pAzo-2** is compatible with imaging techniques and its potential as a cardioselective light-regulated ligand has been validated by the reversible photomodulation of the cardiac rhythm in living zebrafish larvae.

