## Supplemental information for "A photoswitchable ligand targeting β_1_-adrenoceptor enables light-control of the cardiac rhythm"

### Supporting Information

#### SUPPLEMENTARY FIGURES

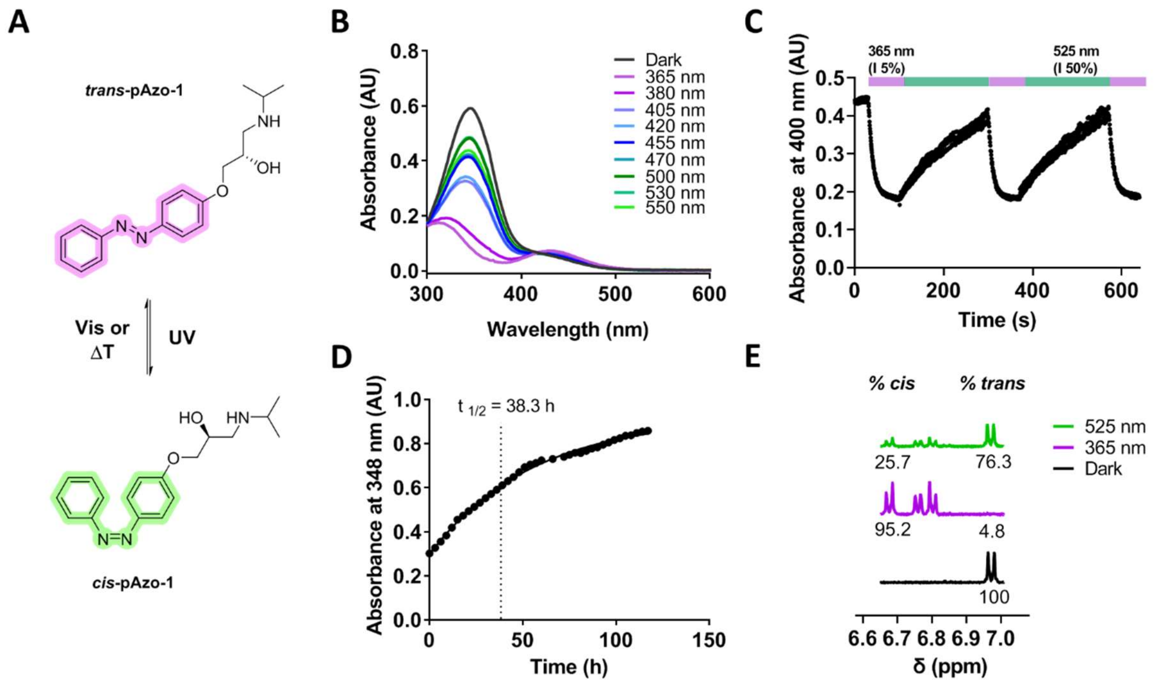

**Figure S1. Photochemical characterization of pAzo-1.** (A) 2D chemical structures of the photoisomers of azobenzene **pAzo-1**. (B) UV-Vis absorption spectra of a 50 μM solution of **pAzo-1** in Epac buffer (0.5% DMSO) under different light conditions. (C) Multiple cis/trans isomerization cycles (365/5350 nm); absorbance was measured at 400 nm. (D) Half-lifetime estimation of *cis*-**pAzo-1** at 25 ºC; absorbance was measured at 348 nm. (E) Photostationary state (PSS) quantification by $^{1}$H-NMR.

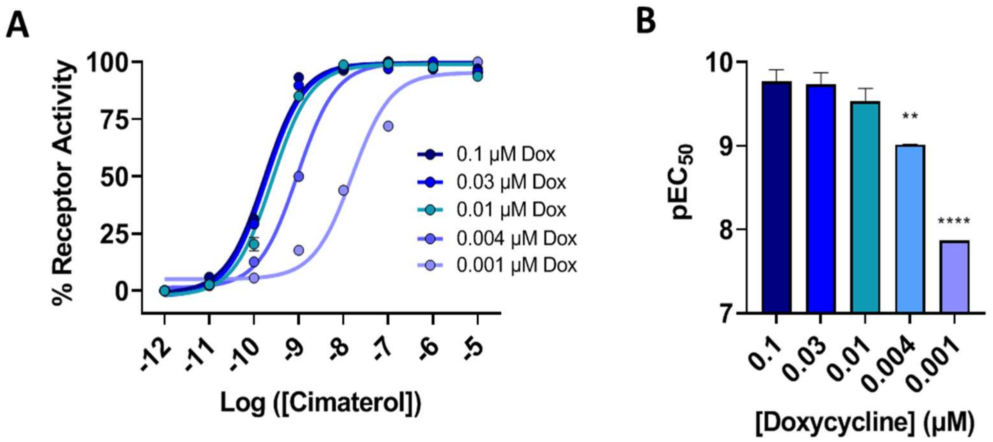

**Figure S2. Optimization of $\beta_1$-AR expression levels in HEK293 iSNAP $\beta_1$-AR cells stably transfected with an Epac cAMP biosensor.** (A) Dose-response curves of cimaterol in cells induced with different levels of doxycycline (0.1-0.001 μM) for 24h. (B) pEC50 values from the cimaterol dose-response curves depicted in graph A. Data are shown as the mean ± SEM of two independent experiments in duplicate. Statistical differences from $pEC_{50}$ values obtained after 0.1 μM doxycycline induction are denoted for adjusted p values as follows: **$p<0.01$ and **$p<0.0001$.

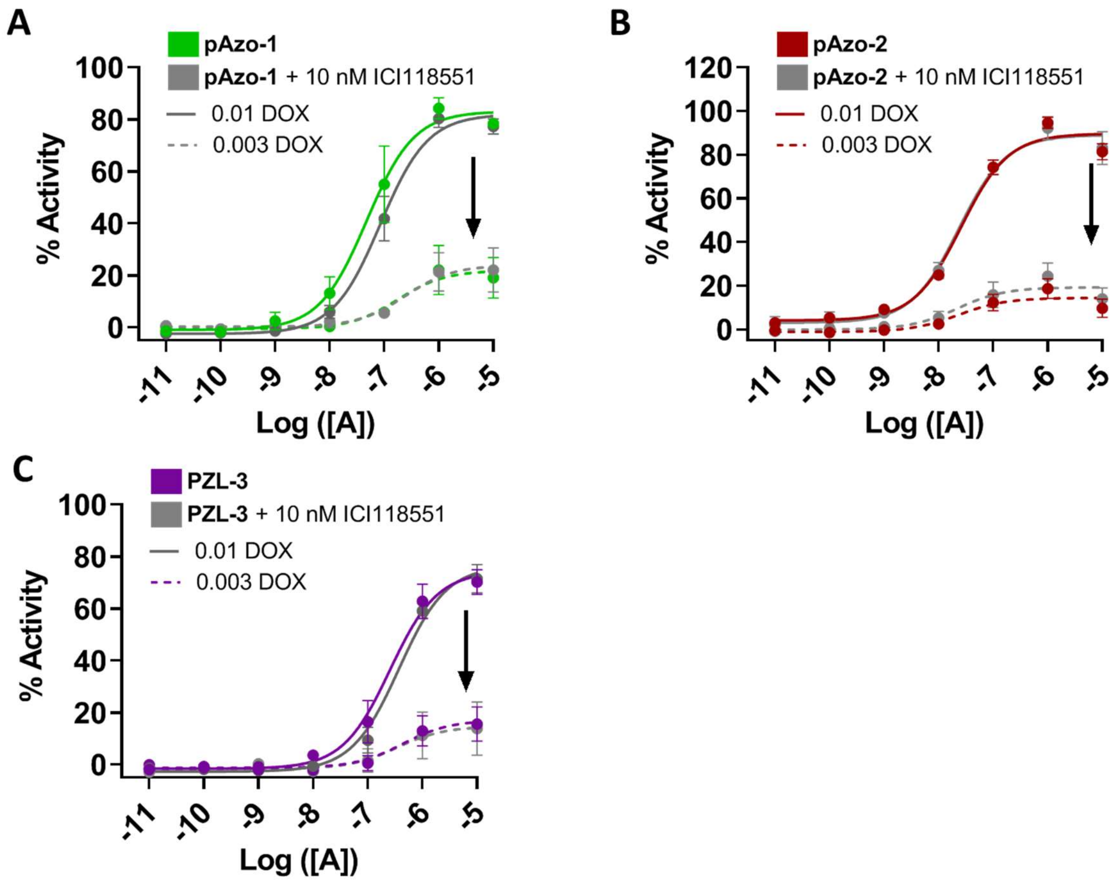

**Figure S3. $\beta_1$-AR functional assays performed in differently induced HEK293 iSNAP $\beta_1$-AR cells stably transfected with an Epac cAMP biosensor.** Dose-response curves of **pAzo-1** (A), **pAzo-2** (B) and **PZL-3** (C) were performed in the presence or absence of a constant concentration of ICI 118551 (10 nM) using cells with different expression levels (24h induction with 0.01 µM dox and 0.003 µM dox respectively). Data are shown as the mean ± S.E.M of four to five independent experiments performed in duplicate.

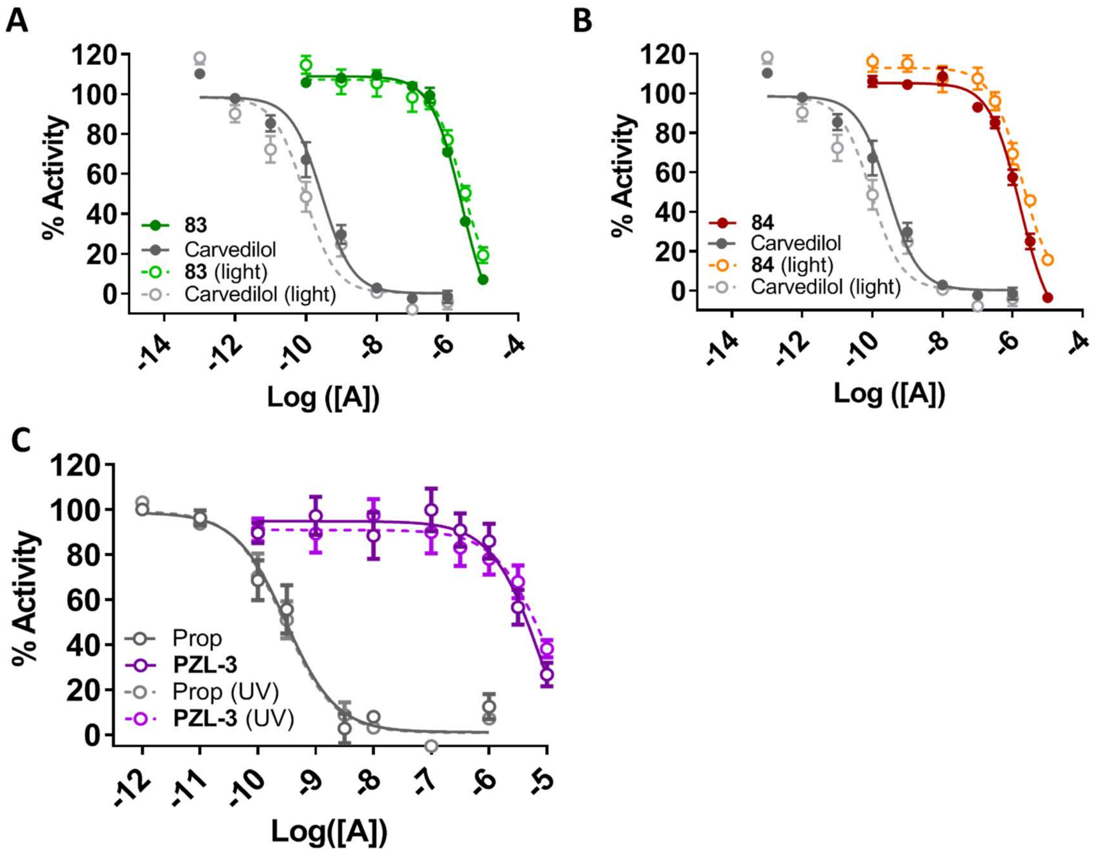

**Figure S4. Light-dependent β$_2$-AR inhibition.** Dose-response curves of **pAzo-1** (A), **pAzo-2** (B), and **pZL-3** (C) performed with a constant concentration of the agonist cimaterol (3 nM) in the dark and under constant violet light (380 nm). Data are shown as the mean ± SEM of four independent experiments in duplicate.

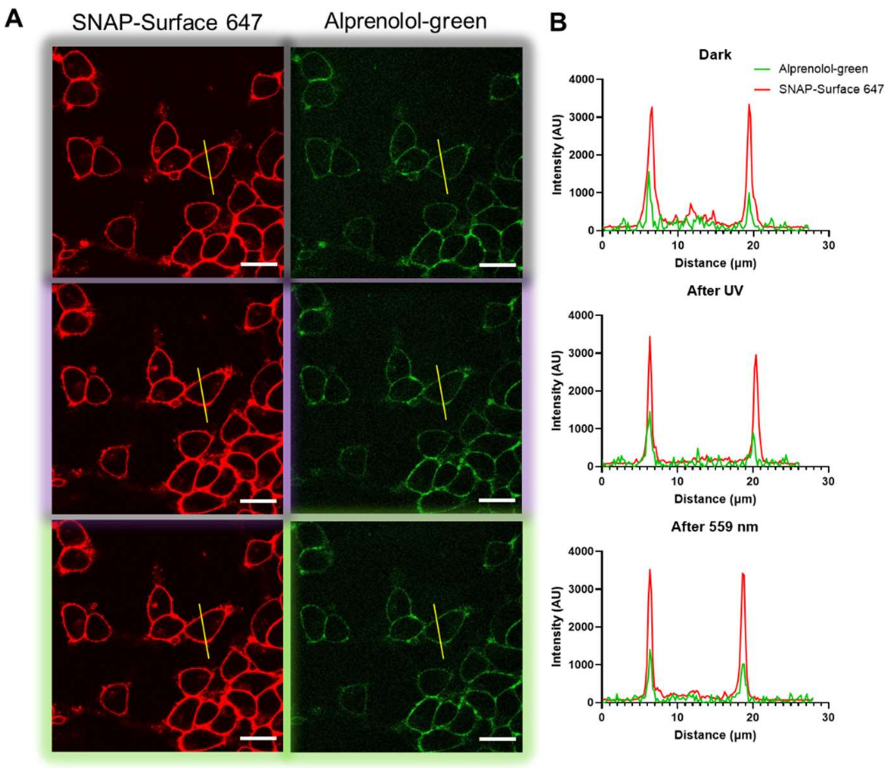

**Figure S5. Light-dependent bleaching control of alprenolol-green binding on $\beta_1$-AR**. (A) Representative confocal fluorescence images of HEK293 iSNAP $\beta_1$AR H188 cells labelled with SNAP-surface-647 and preincubated with alprenolol-green (20 nM) for 1h. Images were obtained in the dark (Top panel), after 10 min violet light exposure (Middle panel) and after 5 min green laser exposure (bottom panel). Scale bars are 20 µm. (B) Plots showing the fluorescence signals measured along the yellow lines shown in (A). The receptor at the plasma membrane is shown in red and the bound alprenolol green is shown in green for the three conditions: dark (top), violet illumination (middle) and green illumination (bottom).

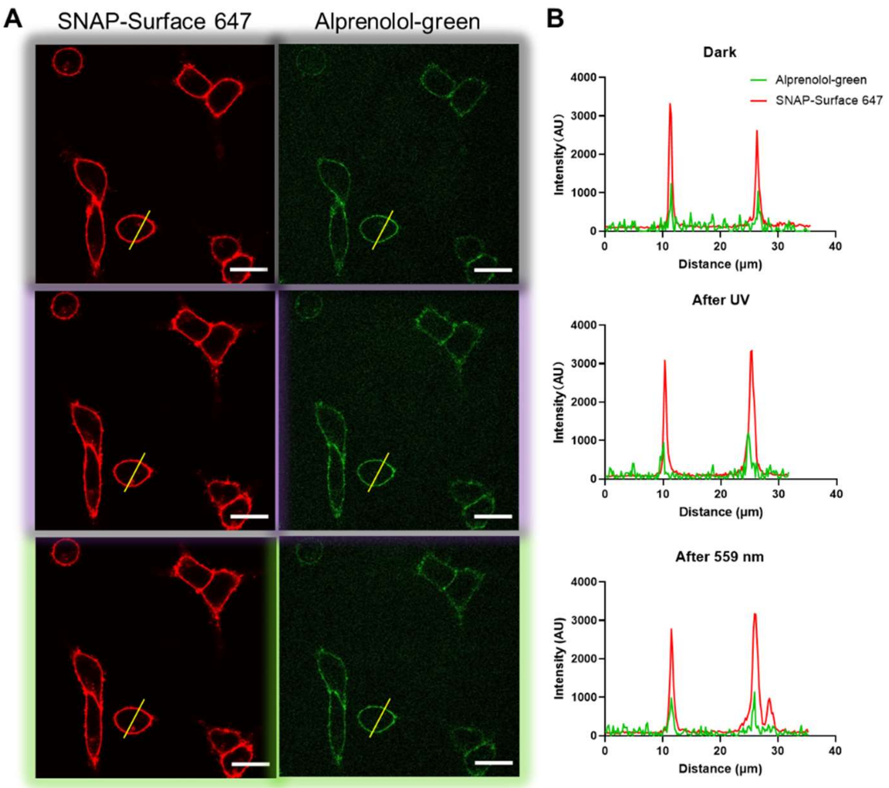

**Figure S6. Light-dependent control of propranolol binding on β$_1$-AR.** (A) Representative confocal fluorescence images of HEK293 iSNAP β$_1$AR H188 cells labelled with SNAP-surface-647 and preincubated with alprenolol-green (20 nM) for 30 min. Cells were treated with 0.1 nM propranolol in dark for 1 h. Images were obtained in the dark (Top panel), after 10 min violet light exposure (Middle panel) and after 5 min green laser exposure (bottom panel). Scale bars are 20 µm. (B) Plots showing the fluorescence signals measured along the yellow lines shown in (A). The receptor at the plasma membrane is shown in red and the bound alprenolol green is shown in green for the three conditions: dark (top), violet illumination (middle) and green illumination (bottom).

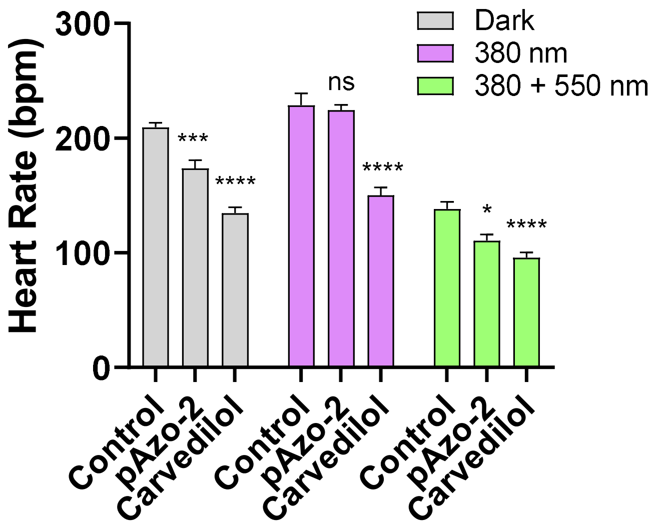

**Figure S7. Optical modulation of the cardiac frequency by *p*-Aazo2.** Raw cardiac frequency of the different experimental groups (Control N=9-18, 25 µM **pAzo-2** N=8-14 and 10 µM carvedilol N=14-15) in the dark, after illumination with a 380 nm light for 1 min and after a subsequent illumination with a 550 nm light. Data are shown as the mean ± SEM of two independent experiments. Statistical differences from control values in each different illumination condition are denoted for adjusted p values as follows: *p<0.05, ***p<0.001, and ****p<0.0001.

#### SUPPLEMENTARY TABLES

##### Table S1. Pharmacological data of photoswitches pAzo-1 and pAzo-2 towards $\beta_2$AR.

| | DARK | | LIGHT | | | |
|---|---|---|---|---|---|---|
| **Cmpd** | **$IC_{50}$ (µM)** | **SEM** | **$IC_{50}$ (µM)** | **SEM** | **PPS[a]** | **SEM** |
| **pAzo-1** | 2.45 | 0.33 | 2.86 | 0.39 | 1.08 | 0.23 |
| **pAzo-2** | 1.65 | 0.26 | 1.78 | 0.18 | 1.17 | 0.33 |
| **PZL-3** | 8.90 | 2.52 | 8.44 | 5.08 | 0.95 | 0.51 |
| **Carvedilol** | 0.00029 | 0.00012 | 0.000086 | 0.00010 | 0.29 | 0.17 |

[a] PPS refers to Photoinduced Potency Shift which is the relation between the measured $IC_{50}$ in light and dark conditions respectively.

##### Table S2. Pharmacological data of pAzo-1, pAzo-2, PZL-3 and cimaterol towards $\beta_1$AR in cellular systems induced two different concentrations of doxycycline.

| | Induction w/ 0.01 µM dox. | | | | Induction w/ 0.003 µM dox | | | |
|---|---|---|---|---|---|---|---|---|
| **Cmpd** | **$EC_{50}$ (nM)** | **SEM** | **Emax (%)** | **SEM** | **$EC_{50}$ (nM)** | **SEM** | **Emax (%)** | **SEM** |
| **pAzo-1** | 53.20 | 32.61 | 84.62 | 2.51 | 111.60 | 68.11 | 18.32**** | 9.52 |
| **pAzo-2** | 26.74 | 7.43 | 90.28 | 2.66 | 10.85 | 8.45 | 11.62** | 4.38 |
| **PZL-3** | 256.08 | 87.40 | 74.37 | 5.73 | 608.21 | 1893.74 | 10.48** | 9.55 |
| **Cimaterol** | 0.36 | 0.05 | 99.98 | 0.02 | 5.28** | 1.39 | 105.41 | 4.62 |

Statistical differences between EC50 and Emax values from the two induction conditions are denoted for adjusted p values as follows: **p<0.01 and ****p<0.0001.

### MATERIAL AND METHODS

#### General Synthetic Methods

All starting materials were obtained from commercial sources and used without further purification unless otherwise stated. Anhydrous solvents were obtained from a solvent purification system (*PureSolv-EN$^{TM}$*) and kept under a nitrogen atmosphere. Butanone was dried over activated molecular sieves prior to use. Reactions were monitored by thin layer chromatography (TLC) on silica gel (60F, 0.2 mm, ALUGRAM Sil G/UV254 *Macherey-Nagel*) and visualized with 254 nm UV light. Flash column chromatography was carried using silica-gel 60 (*Panreac*, 40-63 μm mesh) or by means of a *Biotage* Isolera One automated system with *Biotage* SNAP columns. Analytical HPLC was performed on a *Thermo Ultimate 3000SD* (*Thermo Scientific Dionex*) coupled to a PDA detector and Mass Spectrometer *LTQ XL ESI-ion trap* (*Thermo Scientific)* (HPLC-PDA-MS)) or on a *Waters 2795 Alliance* coupled to a DAD detector (*Agilent 1100*) and an *ESI Quattro Micro* MS detector (*Waters*); HPLC columns used were *ZORBAX Eclipse Plus C18* (4.6x150mm; 3.5μm) and *ZORBAX Extend-C18* (2.1 x 50 mm, 3.5 μm) respectively. HPLC purity was determined using the following binary solvent system: 5% acetonitrile in 0.05% formic acid for 0.5 minutes, from 5 to 100% acetonitrile in 5 minutes, 100% acetonitrile for 1.5 minutes, from 100 to 5% acetonitrile in 2 minutes and 5% acetonitrile for 2 minutes. The flow rate was 0.5 mL/min, column temperature was fixed to 35 ºC and wavelengths from 210-600 nm were registered. The purity of all compounds was determined to be > 95%. Nuclear Magnetic Resonance (NMR) spectroscopy was performed using a *Varian-Mercury 400 MHz* spectrometer. Chemical shifts are reported in δ (ppm) relative to an internal standard (non-deuterated solvent signal). The following abbreviations have been used to designate multiplicities: s=singlet, d=doublet, t=triplet, m=multiplet, br=broad signal, dd=doublet of doublet. Coupling constants (*J*) are reported in Hz. High-resolution mass spectra (HRMS) and elemental composition were performed on a FIA (Flux Injected Analysis) with Ultrahigh-Performance Liquid Chromatography (UPLC) *Aquity (Waters)* coupled to LCT Premier Orthogonal Accelerated Time of Flight Mass Spectrometer (TOF) (*Waters*). Data from mass spectra was analyzed by electrospray ionization in positive and negative mode using MassLynx 4.1 Software (*Waters*). Spectra were scanned between 50 and 1500 Da with values every 0.2 seconds and peaks are reported as *m/z*. IR spectra were recorded neat using *Thermo Nicolet Avatar 360 FT-IR* Spectrometer. Melting points were measured with *Melting Point B-545 (Büchi)*. Optical rotation values were measured in a *Perkin-Elmer 341* polarimeter with the indicated solvents. $[\alpha]_D$ values are reported in degrees and calculated as $c$ x 100 / ($d$ x $m$) where $c$ is the concentration of the sample in g/100 mL, $d$ is the optical way in dm and $m$ is the measured value (mean of 5 measurements). Chiral analytical HPLC was performed on a *Thermo Ultimate 3000SD* (*Thermo Scientific Dionex*) coupled to a *Dionex VWD-3400-RS* detector (*Thermo Scientific*; λ = 362 nm). As chiral HPLC column, a Phenomenex Lux Amylose-2 (4.6 x 250 mm, 5 μM) was used. Enantiomeric excess was determined using the following binary systems: (S1) 0.5% isopropanol + 0.1 % diethylamine + 0.1 % formic acid in acetonitrile, isocratic; 0.6 mL/min.

#### Photophysical Characterization

##### Selection of Photoisomerization Wavelengths

UV-Vis absorbance spectra were recorded using the Spark microplate reader. All samples were prepared with 50 µM of compound in 0.5 % DMSO in aqueous buffer. Samples were measured between 600 nm and 300 nm with 2 nm fixed intervals in 96-well transparent plates (200 µL of compound solution/well). Illumination at the different wavelengths was achieved using the CoolLED pE-4000 light source, set at 50 % intensity. The liquid light guide accessory was pointed directly towards the well containing the studied sample for 3 minutes in continuous mode.

##### Reversibility and repeatability of photoisomerization

Continuous measurements of *trans/cis* isomerization cycles were performed using the Evolution 350 spectrophotometer. Absorbance of the samples (50 µM, 0.5 % DMSO in aqueous buffer) contained in a cuvette was continuously measured at a fixed wavelength in the dark and while illuminated (365 nm or 525 nm). Several isomerization cycles were performed, and transition half-times (τ) were calculated for each compound by plotting absorbance versus time and by fitting the obtained curve to an exponential decay function using GraphPad prism version 8.1.2 (San Diego, CA).

##### Kinetics of *cis* to *trans* isomerization

Thermal relaxation studies were performed at 25ºC by prolonged absorbance measuring (in dark conditions) after samples (50 µM, 0.5 % DMSO in aqueous buffer) had been subjected to 3 min illuminations at 365 nm. Samples were illuminated using the CoolLED light system and measurements were collected using the Spark microplate reader. The relaxation half-life of the compounds was calculated by plotting absorbance readings at a fixed wavelength versus time and by fitting the obtained curve to an exponential decay function using GraphPad prism version 8.1.2 (San Diego, CA). The measured wavelength corresponded to the point where maximum difference in absorbance was observed for *trans* and *cis* isomers.

##### $^1$H-NMR Photostationary State Determination

Data was acquired using a Bruker Avance-III 500 MHz spectrometer equipped with a zaxis pulsed-field gradient triple-resonance (1 H, 13C, 15N) TCI cryoprobe. 1D 1 H spectra were acquired at 285 K with 32 scans using the pulse sequence zgesgppe (water signal suppressed using excitation sculpting and perfect echo) extracted from the Bruker library. Every sample was locked, tuned, and shimmed prior to acquisition. Samples were prepared from a concentrated stock solution of **pAzo-1** and **pAzo-2** (10 mM in $d_6$-DMSO) by dilution in deuterated water (100 µM final). $^1$H NMR spectra of the compounds in dark conditions were initially recorded. Samples were thereafter illuminated for 3 min in the NMR tube using a 96-well LED array plate with the optimal *trans→cis* isomerization wavelength (365 nm). $^1$H NMR spectra were immediately recorded. The sample contained in the NMR tube was finally illuminated at 525 nm to trigger back-isomerization to the *trans* isomer and $^1$H NMR spectra were again collected.

#### *In vitro* Functional Assays

##### Cell Culture and Transfection

To evaluate the activity of the ligands towards β-AR, two different cellular systems were employed. The activity of photoswitchable ligands against $\beta_2$-AR was evaluated using HEK293 H188 M1 cells, which stably express a cAMP Epac-$S^{H188}$ FRET biosensor[1] and was reported in our previous paper.[2] On the other hand, to evaluate the functional activity of $\beta_1$-AR, a iSNAP $\beta_1$-AR HEK-293 polyclonal cell line were used, which were established by co-transfecting Flp-In™ T-REx™ host cell line (Invitrogen) with an excess of pOG44 plasmid encoding Flp recombinase and pcDNA™5/FRT/TO/SNAP-β1AR expression vectors. These cells can express $\beta_1$-AR upon induction with doxycycline and were used to establish a new polyclonal stable cell line with the cAMP Epac-$S^{H188}$ FRET biosensor[1] following the transfection and selection protocols previously described.[2] We have named this cell line HEK293 iSNAP $\beta_1$AR H188. We maintained both HEK293 cell lines stably expressing the Epac-$SH^{188}$ cAMP biosensor at 37ºC, 5% $CO_2$ in 4,5 g/L D-glucose Dulbecco's Modified Eagle Medium (DMEM, GIBCO) supplied with 10% heat inactivated FBS (GIBCO) and 1% penicillin-streptomycin (10,000 U/mL, GIBCO). In the case of HEK293 iSNAP $\beta_1$AR H188 cells, the medium was additionally supplemented with Hygromycin B (100 µg/mL) and Blasticidin (15 µg/mL) since this cell line is also stable for the expression of the human ADRB1. All cell lines were split when reaching 75-90% confluence and detached by trypsin digestion.

##### General Methods

All assays were performed at room temperature. Adherent cells were grown in T-175 flasks or 150-mm dishes to 75-90% confluence. Cells were detached by rinsing once with PBS (GIBCO), followed by incubation with Trypsin-EDTA (Sigma-Aldrich) for 5 minutes until detachment of cells was observed. Cells were then centrifugated; in parallel, 10 µL of the single cell suspension were counted using a Neubauer Chamber. The supernatant was carefully removed, and cells were resuspended in DMEM complete medium to obtain a cell solution with 1.0 x $10^6$ cells/mL. 100,000 cells per well were seeded in a transparent 96-well microplate (Thermo Scientific Nunc Microwell) and left at 37ºC with 5% $CO_2$ for approximately 24h. In the case of assays performed using HEK293 iSNAP $\beta_1$AR H188 receptor was induced by addition of 0.01 µg/mL doxycycline to the medium for 18-24 h. cAMP EPAC sensor buffer (14 mM NaCl, 50 nM KCl, 10 nM $MgCl_2$, 10 nM $CaCl_2$, 1 mM HEPES, 1.82 mg/mL Glucose, pH 7.2) was used as the assay medium in all FRET-based experiments. In assays performed using the HEK293 H188 M1 cell line, cAMP EPAC sensor buffer was supplemented with 100 µM IBMX. Fluorescence values were measured using a Tecan Spark M20 multimode microplate reader equipped with the Fluorescence Top Standard Module with defined wavelength settings (excitation filter 430/20 nm and emission filters 485/20 nm and 535/25 nm). FRET ratio was calculated as the ratio of the donor emission ($td^{cp173}V$, 485 nm) divided by the acceptor emission (mTurq2Δ, 535 nm). The FRET ratio was normalized to the effect of the buffer (0%) and the maximum response obtained with cimaterol (100%). External light was applied using the 96-well LED array plate (LEDA Teleopto). Each set of experiments was performed three to five times with each concentration in duplicate or triplicate.

Dose-Response Assays

To perform cimaterol-stimulated assays with the azobenzene ligands in HEK293 H188 M1 cells we prepared two different plates, one for each light condition. For all assays, both plates were left to incubate with the studied compounds for 45 minutes at room temperature. To induce photoswitching, the "light plate" was exposed to continuous illumination (380 nm) during the incubation time using the LED array plate (LEDA Teleopto). Fluorescence values were thereafter measured for 30 minutes. Dose-response curves for each compound were obtained using a constant concentration of the agonist cimaterol (3.16 nM). In order to evaluate the activity of the developed azobenzenes in the HEK293 iSNAP $\beta_1$AR H188 cell line, only one plate was used to evaluate both light conditions. Different concentrations of the tested compounds were incubated with the cells for 45 min in the dark at room temperature. Fluorescence values were thereafter measured. The same plate was then continuously illuminated for 45 min (380/405 nm) using the Teleopto LED array system. Fluorescence values were measured again.

Real-Time Assays

Assays to assess the dynamic control of receptor activity were carried out using a constant concentration of the tested azobenzene. Response of the cells with buffer and buffer supplemented with cimaterol was also evaluated. Cells were left to incubate with the compounds for 1h and fluorescence was measured. Then, the plate was continuously illuminated with light at 380 nm for 10 minutes and fluorescence values were recorded. Immediately after, the plate was illuminated in continuous mode for 10 minutes with light at 550 nm and fluorescence was measured. An additional light cycle was applied to ensure the reversibility in receptor activity triggered by light was reproducible over time.

**Confocal microscopy assays**

Cell Culture

Polyclonal inducible HEK293 iSNAP $\beta_1$AR H188 cells were grown at 37°C, 5% $CO_2$ and >95% humidity in DMEM/F-12 (Fisher Scientific) with GlutaMax™. Culture medium was supplemented with 10% Fetal Bovine Serum (InVitro), 100 µg/ml hygromycin B (Fisher Scientific), 15 µg/ml blasticidin (Fisher Scientific), and Phenol Red (Gibco). The day before imaging, cells were induced with 1 µg/ml tetracycline (abcr) and seeded onto 8 well chambered coverslips (Ibidi) in culture medium (30,000 cells/well). On the second day of the experiment, cells were pre-labelled for 30 min with 2.5 µM SNAP-Surface 647 (New England BioLabs) in imaging medium (DMEM/F12 without Phenol Red, Life Technologies) at room temperature (RT). After labelling, cells were washed and maintained in imaging medium. Cells were then incubated with 20 nM alprenolol-green (Cisbio, PerkinElmer) in binding buffer (NaCl, KCl, $MgCl_2 \cdot 6H_2O$, $CaCl_2$, HEPES and Glucose) (Sigma Aldrich) for 30 min. The competitive ligand, either 0.1 nM Propranolol (Sigma Aldrich) or 100 nM **pAzo-2**, was finally added, and samples were imaged after 60 min of incubation in dark conditions at RT. To perform competitive binding assays, cells were preincubated with 50 nM of alprenolol-green (Cisbio, PerkinElmer) in binding buffer for 30 min, and then treated with different concentrations of propranolol for 60 min in dark conditions.

Fluorescence imaging and image analysis

In both cases, live cell imaging was performed with an ***Olympus IX81 confocal microscope***. Alprenolol-green was excited by a 488 nm laser and detected from 500 to 600 nm. SNAP-Surface 647 was excited at 635 nm and detected from 650 to 750 nm. Images were acquired at the same position before and after 10 min illumination with 380 nm UV light (11.12 mW/cm$^2$) and after 5 min illumination with a 559 nm laser (2.64 mW/cm$^2$).

Intensity profiles were measured using FIJI, whereas the signal intensity at the membrane was measured as previously[3] but, here, with the use of an automated script in MATLAB. Each data point is a collection of data from three independent experiments, and error bars represent the standard error of the mean. About 2000 cells were analyzed in each competitive binding assay.

**Cardiac frequency monitorization in zebrafish larvae**

Fish Husbandry and Larvae Production

Adult wild-type zebrafish were purchased from EXOPET (Madrid, Spain) and maintained in fish water (reverse-osmosis purified water containing 90 µg/mL of Instant Ocean (Aquarium Systems) and 0.58 mM $CaSO_4 \cdot 2H_2O$) at 28 ± 1°C in the Research and Development Center of the Spanish Research Council (CID-CSIC) facilities under standard conditions. Embryos were obtained by in-tank group breeding with a 3:2, female:male ratio per tank. Embryos were collected and maintained in 500 mL glass containers at 1 individual/mL density in fish water at 28.5 °C on a 12 light:12 dark photoperiod. Larvae were not fed before or during the experimental period. All procedures were approved by the Institutional Animal Care and Use Committees at the CID-CSIC and conducted in accordance with the institutional guidelines under a license from the local government (agreement number 9027).

General Methods

7 days post-fertilization (dpf) zebrafish larvae were observed under a Motic SMZ-171 dissecting microscope fitted with a GigE camera (iDS - Imaging Development Systems GmbH). Cardiac rhythms were measured in dark conditions and light from 7 to 10 larvae from each treatment and light condition. Larvae were placed over 3% methylcellulose with a drop of medium water and positioned to view the heart. Videos were recorded in dark conditions for 30 seconds using uEye Cockpit application (iDS software) at a frame rate of 40 fps. To prevent light contamination from the microscope, a red filter was placed between the light source and the examined animals. Videos were analyzed with a MATLAB algorithm (version R2010b, MathWorks, USA) which provided the cardiac rhythm (heart beats per minute) for each individual. The program was based on a code to measure the heart rate from a video of a human fingertip captured with a smartphone camera (GitHub: uavster/Video2HeartRate). Briefly, after the video is recorded the brightness signal of the heart region is computed as pixel variations between frames. Then, a band-pass filter is applied (Butterworth filter) to attenuate all frequencies outside the interest band and a Fourier transform is calculated using the Fast Fourier Transform (FFT) algorithm to translate the signal from the time domain to the frequency domain.

Finally, the maximum FFT amplitude is obtained, and the corresponding frequency transformed to beats per minute (BPM).

Light-dependent modulation of the cardiac rhythm *in vivo*

To assess differential cardiac modulation enabled by the two photoisomers of **pAzo-2**, zebrafish larvae (7 dpf) were exposed to the different experimental conditions for 1.5 to 2 h, at 28.5°C (POL-EKO APARATURA Climatic chamber KK350). The different experimental groups, control (1 % DMSO), 10 µM carvedilol and 25 µM of **pAzo-2** were kept in 6-well plates for the incubation (15-20 individuals per well in 2 mL of aqueous treatment). Two different plates of zebrafish larvae were used to test both, dark and light conditions. The dark experimental group was kept in dark conditions for 1.5 to 2 h and cardiac rhythm was then monitored for each larva. The light plate was incubated in dark conditions for 1 to 1.5 h and illuminated continuously for 1 min using 380 nm light (13.3 V). In order to assess light-dependent effects related to treatment with **pAzo-2**, animals were left to acclimate for 30 min prior to heart monitorization, which avoids non-specific alterations of the cardiac rhythm induced by the visual stimulus. These effects are important with white light and are also observed with violet light, considering that zebrafish have specific violet light photoreceptors.[4] Finally, to evaluate light-triggered back-isomerization, an additional plate was exposed for 1 h in dark conditions. 380 nm (1 min, 13.3 V) light was then applied, and the plate was maintained in dark conditions for 30 min. Finally, the plate was illuminated again at 550 nm (1 min, 13.3 V) and the cardiac rhythm was monitored 30 min after light application.

**Data Analysis**

All experiments were analyzed using GraphPad Prism 8.1.1 (GraphPad Software, San Diego, CA). Data are shown as the mean ± SEM unless otherwise stated. Statistical differences were considered significant when $p < 0.05$ unless otherwise stated. For the analysis of *in vitro* assays, stimulation Dose-Response data was fitted using the log (agonist) vs. response (three parameters) function. Inhibition Dose-Response data was fitted using the log(antagonist) vs. response (three parameters) function. Statistical analysis comparing the $EC_{50}$ or $IC_{50}$ values of the compounds in dark and light conditions was performed by unpaired Student's t-test. Statistical analysis comparing the three light conditions in Real Time Assays was performed by a one-way ANOVA followed by Tukey's multiple comparisons test. Data obtained from fluorescence competitive binding assays were fitted with GraphPad prism 9 (Fitting model - Binding Competitive- One site - Fit logIC50).

$$Y = Bottom + \frac{Top - Bottom}{1 + 10^{(X - LogIC5\ )}}$$

Where Top and Bottom are plateaus in the units of Y axis., X is the log of concentration of unlabelled ligand (**pAzo-2** or propranolol). Y is the intensity of bound alprenolol-green and LogIC50 is the log of the concentration of competitor (**pAzo-2** or propranolol) that results in binding half-way between Bottom and Top. For the analysis of the *in vivo* experiments, statistical differences were assessed by performing a one-way ANOVA followed by Tukey's multiple comparisons test.

#### COMPOUND SYNTHESIS AND CHARACTERIZATION

##### *(E)*-4-((4-Methoxyphenyl)diazenyl)phenol (3)

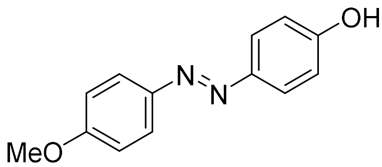

4-Methoxyaniline (**1**) (500 mg, 4.1 mmol, 1.0 eq) was taken up in water (4 mL) and cooled down in an ice bath. HCl (2.1 mL) was then added slowly and the solution was stirred for 10 min at 0ºC.

A solution of sodium nitrite (630 mg, 9.1 mmol, 2.25 eq) in water (6 mL) was then added dropwise and the mixture was left to stir for 10 more minutes in an ice bath. The resulting red slurry was finally added slowly on top of an ice-cold solution of phenol (764 mg, 8.1 mmol, 2.0 eq) in NaOH 10% (8 mL) and the reaction was left to stir for 1 h. A brown precipitate had been formed. The suspension was filtered off with cold water and dried to yield 730 mg of **3** (yield = 79 %), m.p. 139.5 - 139.6 ºC.

**$^1$H NMR** (400 MHz, Chloroform-*d*) δ = 7.89 (d, *J* = 8.8 Hz, 2H), 7.84 (d, *J* = 8.8 Hz, 2H), 7.00 (d, *J* = 8.8 Hz, 2H), 6.94 (d, *J* = 8.8 Hz, 2H), 3.89 (s, 3H). Spectroscopic data is in agreement with that reported in the literature.[5]

##### *(S,E)*-1-(4-(Oxiran-2-ylmethoxy)phenyl)-2-phenyldiazene (4)

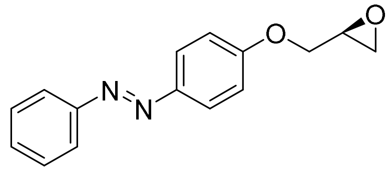

(*E*)-4-(Phenyldiazenyl)phenol (**2**) (200 mg, 1.01 mmol, 1.0 eq) was taken up in butanone (3 mL) in a nitrogen atmosphere. Potassium carbonate (418 mg, 3.03 mmol, 3.0 eq) was added and the reaction was left to stir for 10 minutes. (*R*)-epichlorohydrin (0.396 ml, 5.04 mmol, 5 eq) was finally added and the reaction was heated to reflux and left to stir overnight. The reaction mixture was filtered, and the precipitate washed with acetone (10 mL x 3). The resulting solution was dried under reduced pressure to yield a brown oil. The crude product was further purified by column chromatography (hexane/EtOAc 3:1). Oxirane **4** was isolated as an orange solid (703 mg, 90% yield), m.p. 85 - 87 ºC.

**$^1$H NMR** (400 MHz, Chloroform-*d*) δ = 7.92 (d, *J* = 9.1 Hz, 2H), 7.88 (dd, *J* = 7.2, 2 Hz, 2H), 7.55-7.48 (m, 2H), 7.48-7.39 (m, 1H), 7.04 (d, *J* = 9.1 Hz, 2H), 4.32 (dd, *J* = 11.0, 3.0 Hz, 1H), 4.03 (dd, *J* = 11.0, 5.7 Hz, 1H), 3.40 (m, 1H), 2.94 (dd, *J* = 4.8, 4.0 Hz, 1H), 2.79 (dd, *J* = 4.9, 2.6 Hz, 1H). **$^{13}$C NMR** (101 MHz, Chloroform-*d*) δ = 161.0, 152.8, 147.4, 130.6, 129.2, 124.9, 122.7, 115.0, 69.1, 50.1, 44.8. **IR** (neat): *υ* = 3066, 3003, 2919, 2874, 1603, 1580, 1497, 1242, 1142, 1027, 835, 764, 689. **HRMS** (ESI +): *m/z* calcd for $C_{15}H_{15}N_2O_2^+$[M + H]$^+$ = 255.1134; found 255.1121. **$[\alpha]^{25}_D$**= + 7.6 (c = 1.6, MeOH). (*R*)-**81** (981 mg, 96 %) was produced following the same protocol but using (*S*)-epichlorohydrin (1.6 mL, 15.7 mmol).

***(S,E)*-1-(4-Methoxyphenyl)-2-(4-(oxiran-2-ylmethoxy)phenyl)diazene (5)**

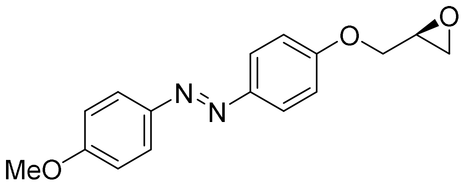

(*E*)-4-((4-Methoxyphenyl)diazenyl)phenol (**3**) (630 mg, 2.8 mmol, 1.0 eq) was taken up in butanone (6 mL). Potassium carbonate (1.1 g, 8.3 mmol, 3.0 eq) was then added to the solution and the mixture was left to stir under nitrogen for 10 min. (*R*)-epichlorohydrin (1.08 mL, 13.8 mmol, 5.0 eq) was then added and the reaction was heated to reflux and left to stir overnight. The mixture was filtered, and the precipitate washed with acetone (3 x 20 mL). The obtained red solution was evaporated under reduced pressure to yield **5** (703 mg, 2.47 mmol, 90 % yield) as a brown solid, m.p. 130.3 – 131.6 °C.

**$^{1}$H NMR** (400 MHz, Chloroform-*d*) δ = 7.88 (d, *J* = 8.8 Hz, 2H), 7.87 (d, *J* = 8.8 Hz, 2H), 7.02 (d, *J* = 7.6 Hz, 2H), 7.00 (d, *J* = 7.6 Hz, 2H), 4.31 (dd, *J* = 11.0, 2.6 Hz, 1H), 4.03 (dd, *J* = 11.0, 5.8 Hz, 1H), 3.89 (s, 3H), 3.39 (ddt, *J* = 5.8, 4.2, 2.6 Hz, 1H), 2.94 (dd, *J* = 4.9, 4.2 Hz, 1H), 2.79 (dd, *J* = 4.9, 2.6 Hz, 1H). **$^{13}$C NMR** (101 MHz, Chloroform-*d*) δ= 161.8, 160.5, 147.5, 147.2, 124.5, 124.5, 115.0, 114.3, 69.1, 55.7, 50.2, 44.8. **IR** (neat): *υ* = 3096, 3076, 3014, 2928, 2840, 1600, 1580, 1495, 1238, 1148, 1026, 840. **HRMS** (ESI +): *m/z* calcd for $C_{16}H_{17}N_2O_3^+$[M + H]$^+$ = 285.1239; found 285.1332. **$[\alpha]^{25}_{D}$**= + 7.5 (c = 4.6, MeOH).

***(S,E)*-1-(Isopropylamino)-3-(4-(phenyldiazenyl)phenoxy)propan-2-ol (pAzo-1)**

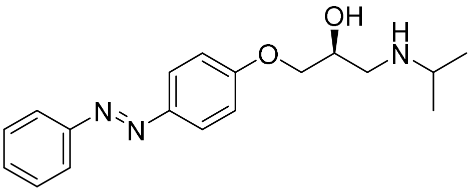

Oxirane **4** (100 mg, 0.39 mmol, 1.0 eq) was dissolved in isopropylamine (1.68 mL) in a tube and heated at 40ºC for 72 h. The reaction mixture was then concentrated under reduced pressure. The isolated crude was purified by column chromatography (10 % MeOH in DCM, 1 % Ammonia) and azobenzene **pAzo-1** was isolated as a yellow solid (30 mg, 24 % yield), m.p. 108.4–109.8 °C.

**$^{1}$H NMR** (400 MHz, Methanol-*$d_4$*) δ = 8.52 (s, 1H), 7.91 (d, *J* = 9.0 Hz, 2H), 7.84 (d, *J* = 9.0, 2H), 7.53–7.41 (m, 3H), 7.12 (d, *J* = 9.0 Hz, 2H), 4.24 (dtd, *J* = 9.6, 5.4, 3.1 Hz, 1H), 4.13 ((dd, *J* = 9.6, 5.4 Hz, 1H), 4.09 (dd, *J* = 8.3, 5.4 Hz, 1H), 3.40 (h, *J* = 6.6 Hz, 1H), 3.25 (dd, *J* = 12.6, 3.1 Hz, 1H), 3.11 (dd, *J* = 12.6, 9.6 Hz, 1H), 1.34 (dd, *J* = 6 .6 Hz, 3H), 1.33 (dd, *J* = 6 .6 Hz, 3H). **$^{13}$C NMR** (101 MHz, Methanol-*$d_4$*) δ = 168.8, 161.1, 152.6, 147.2, 130.3, 128.8, 124.3, 122.1, 114.6, 70.0, 65.7, 50.4, 18.2, 17.7. **IR** (neat): *υ* = 3303, 3070, 2964, 2922, 2870, 1600, 1582, 1501, 1250, 1141, 837, 689. **HRMS** (ESI +): *m/z* calcd for $C_{18}H_{24}N_3O_2^+$[M + H]$^+$ = 314.1869; found 314.1896. **$[\alpha]^{25}_{D}$**= + 10.5 (c = 2.5, MeOH). (*R*)-enantiomer of **pAzo-1** (101 mg, 59 %) was synthesised from (*R*)-**4** (140 mg, 0.55 mmol) as described above. **Chiral HPLC** (system 4): $t_R$ = 14.4 min, ee 93%.

***(S,E)*-1-(Isopropylamino)-3-(4-((4-methoxyphenyl)diazenyl)phenoxy)propan-2-ol (pAzo-2)**

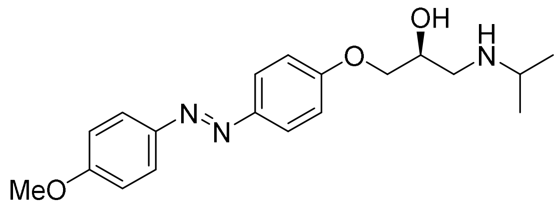

Oxirane **5** (200 mg, 0.70 mmol, 1.0 eq) was dissolved in isopropylamine (3 mL) and the reaction was heated to reflux for 48 h. The reaction mixture was then concentrated *in vacuo*. The crude material was then purified by column chromatography (10% MeOH in DCM, 1 % ammonia) and azobenzene **pAzo-2** isolated as a yellow solid (150 mg, 62.1 % yield), m.p. 137.2–137.6 ºC.

**$^{1}$H NMR** (400 MHz, DMSO-$d_6$) δ = 7.85 (d, *J* = 9.1 Hz, 2H), 7.84 (d, *J* = 9.1 Hz, 2H), 7.12 (d, *J* = 9.1 Hz, 2H), 7.11 (d, *J* = 9.1 Hz, 2H) , 4.08 (dd, *J* = 9.1, 4.1 Hz, 1H), 4.04-3.94 (m, 2H), 3.86 (s, 3H), 2.90 (h, *J* = 6.3 Hz, 1H), 2.83 (dd, *J* = 12.1, 3.7 Hz, 1H), 2.72 (dd, *J* = 12.1, 7.1 Hz, 1H), 1.07 (d, 6.3 Hz, 6H). **$^{13}$C NMR** (101 MHz, DMSO-$d_6$) δ= 161.50, 160.84, 146.21, 124.15, 115.09, 114.57, 70.91, 67.42, 55.63, 48.75, 21.71. **IR** (neat): *υ* = 3316, 2964, 2927, 2835, 1595, 1581, 1495, 1238, 1146, 1025, 841. **HRMS** (ESI +): *m/z* calcd for $C_{19}H_{26}N_3O_3^+$[M + H]$^+$ = 344.1974; found 344.1942. **[α]$^{25}_{D}$**= + 1.72 (c = 2.5, MeOH).

### NMR SPECTRA, PSS& %EE DETERMINATION

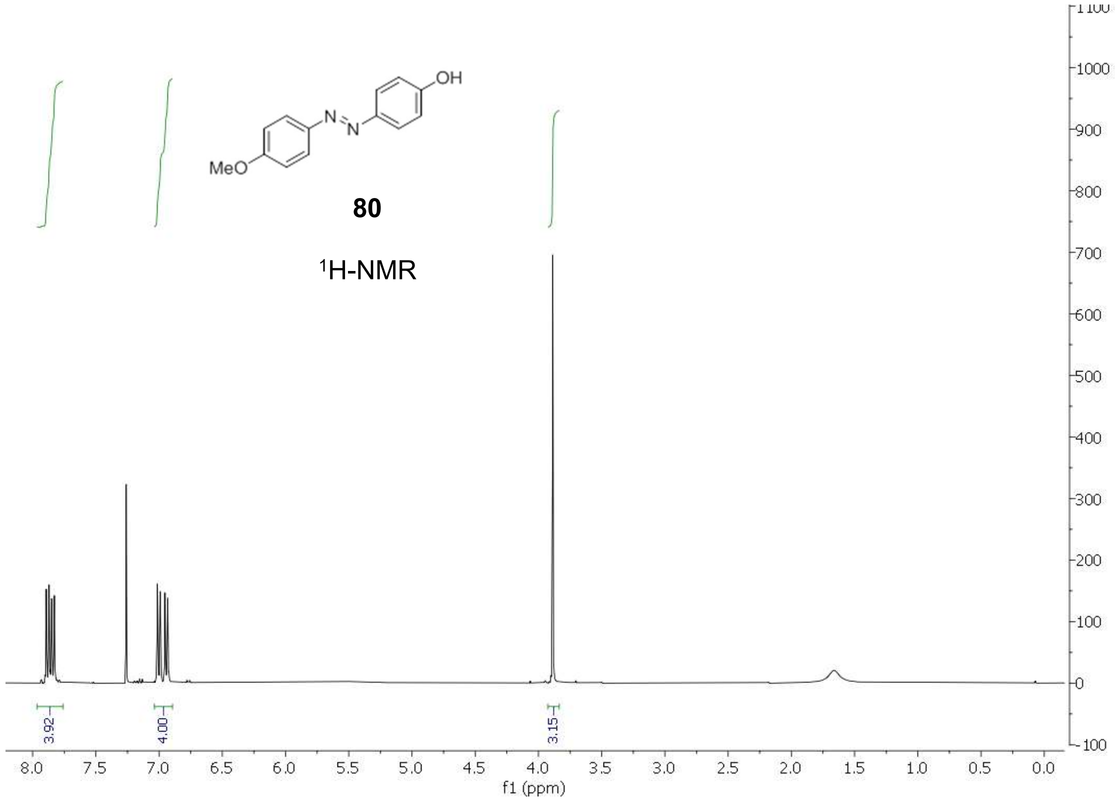

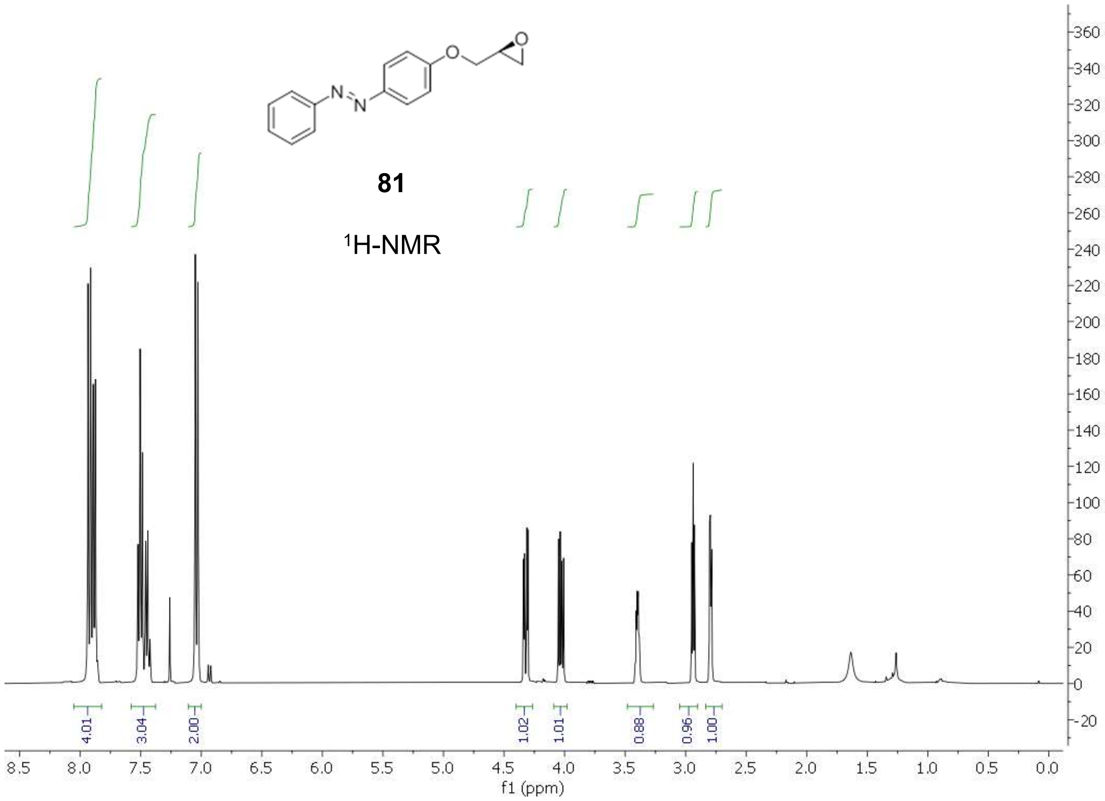
81
1H-NMR
4.01
3.04
2.00
1.02
1.01
0.88
0.96
1.00
f1 (ppm)

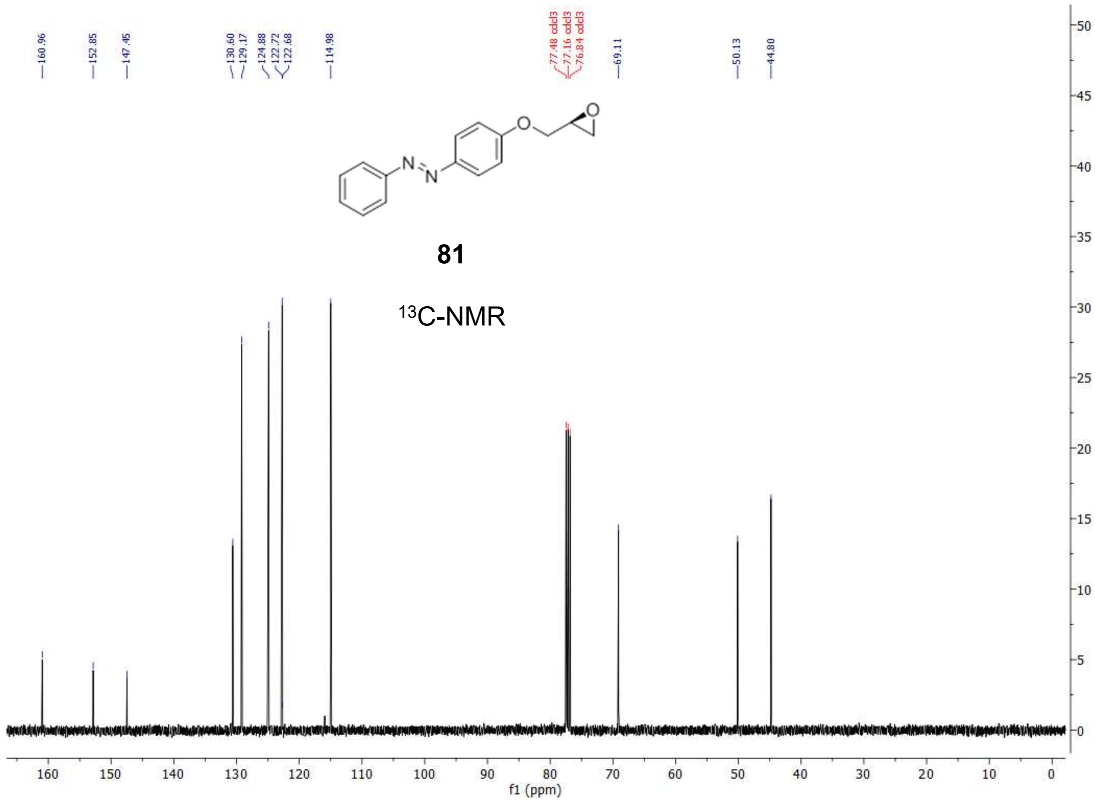
160.96
152.85
147.45
130.60
129.17
124.88
122.72
122.68
114.98
77.48 cdcl3
77.16 cdcl3
76.84 cdcl3
69.11
50.13
44.80
81
13C-NMR
f1 (ppm)

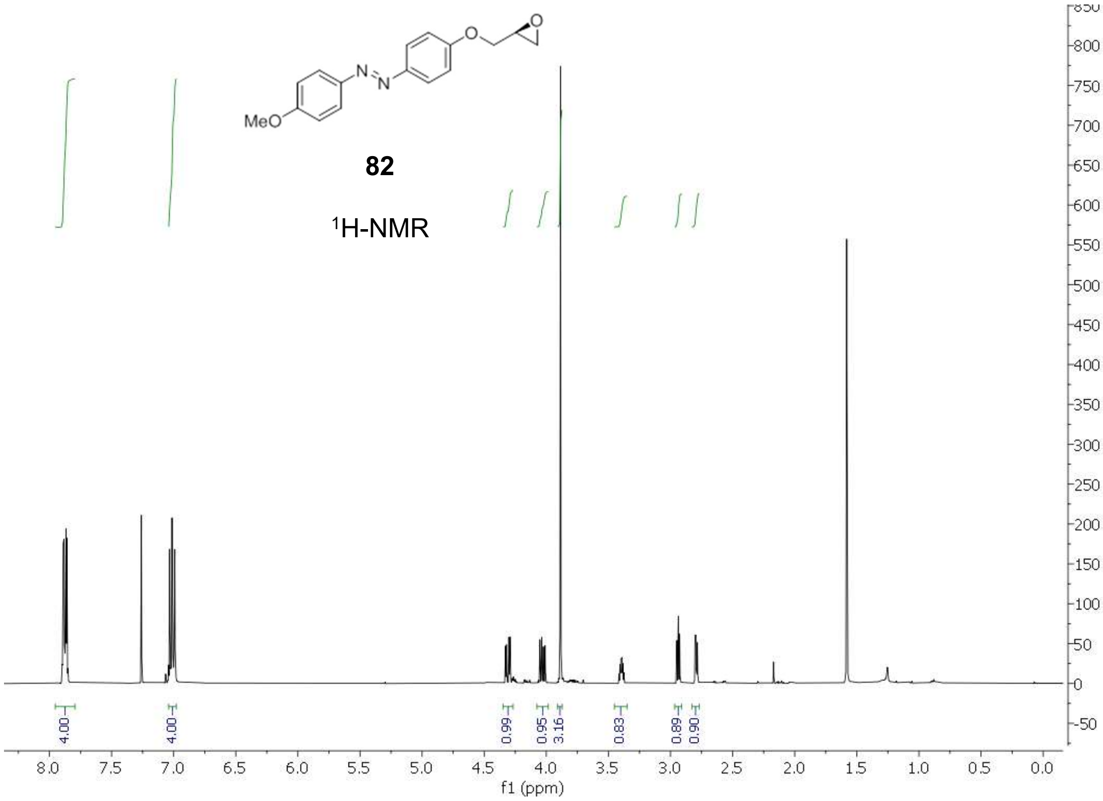
MeO
82
1H-NMR
4.00
4.00
0.99
0.95
3.16
0.83
0.89
0.90
f1 (ppm)

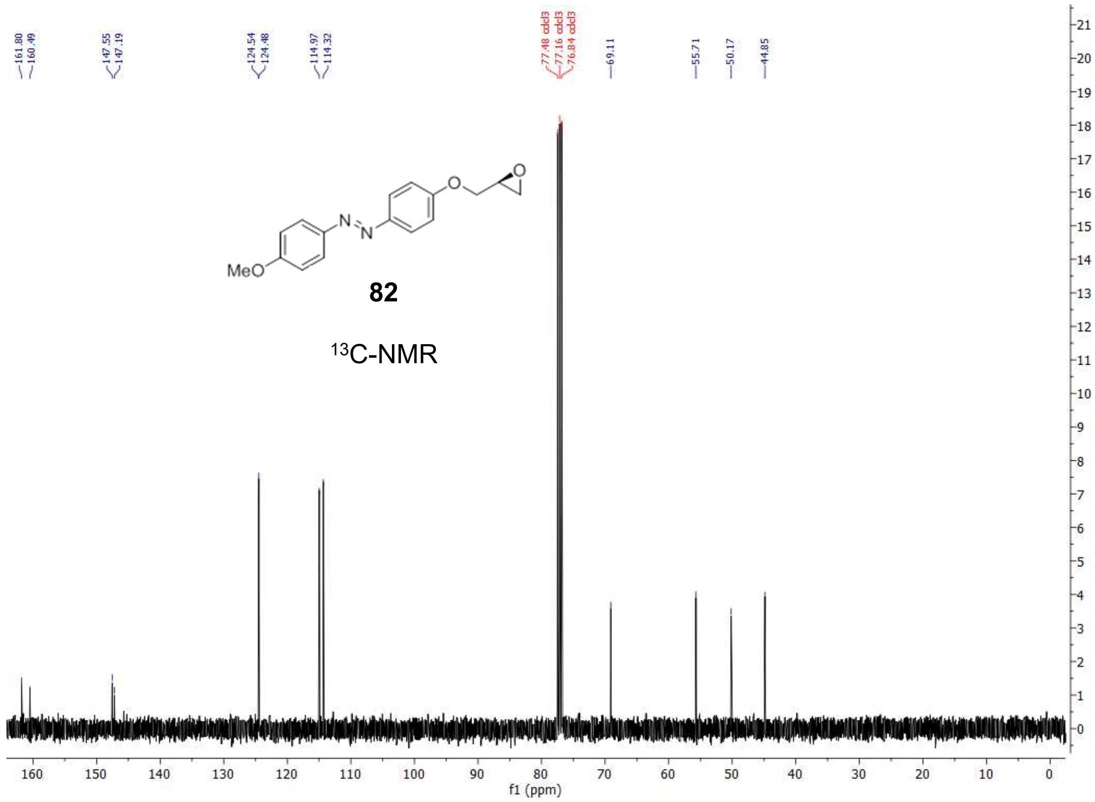
161.80
160.49
147.55
147.19
124.54
124.48
114.97
114.32
77.48 cdcl3
77.16 cdcl3
76.84 cdcl3
69.11
55.71
50.17
44.85
MeO
82
13C-NMR
f1 (ppm)

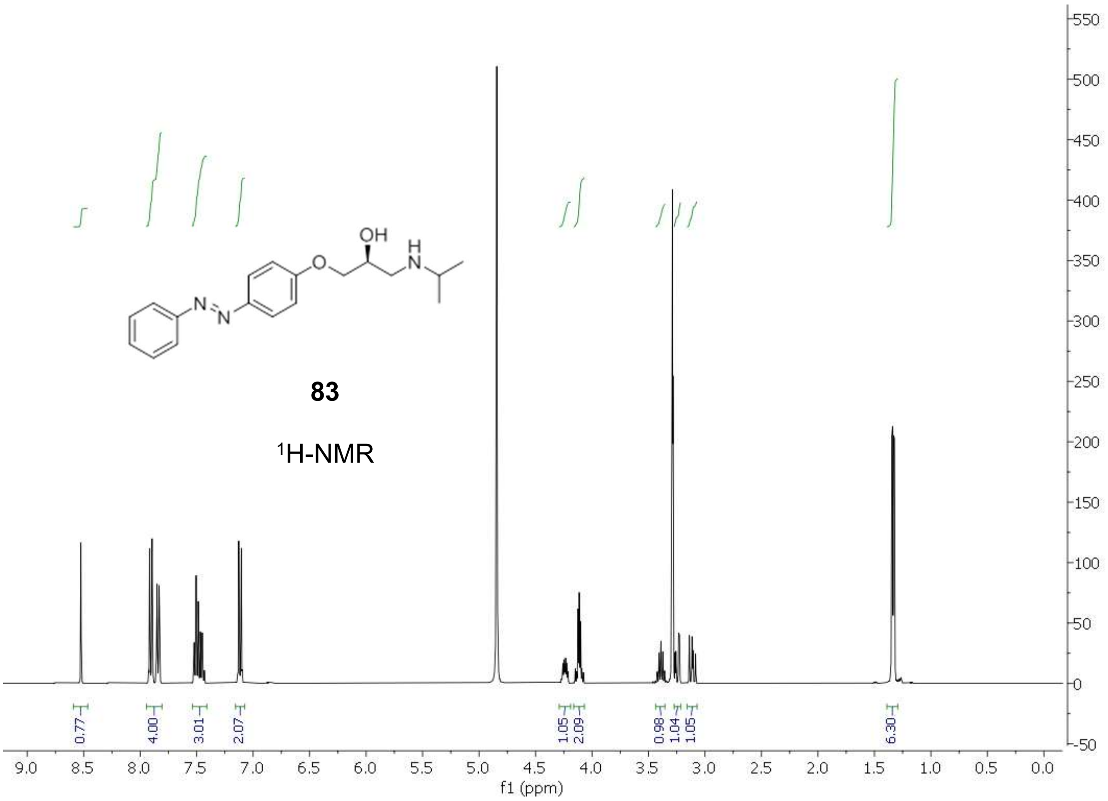
OH
H
N
N
N
O
83
1H-NMR
0.77
4.00
3.01
2.07
1.05
2.09
0.98
1.04
1.05
6.30
f1 (ppm)

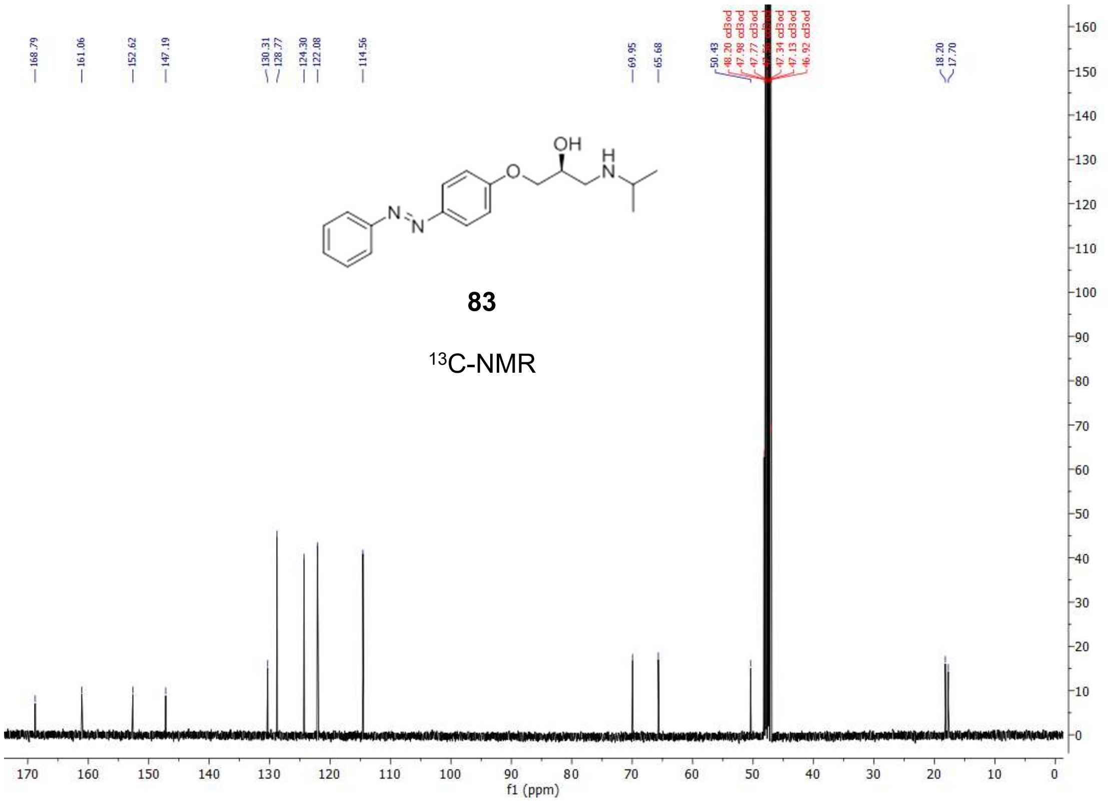
168.79
161.06
152.62
147.19
130.31
128.77
124.30
122.08
114.56
69.95
65.68
50.43
18.20
17.70
OH
H
N
N
N
O
83
13C-NMR
f1 (ppm)

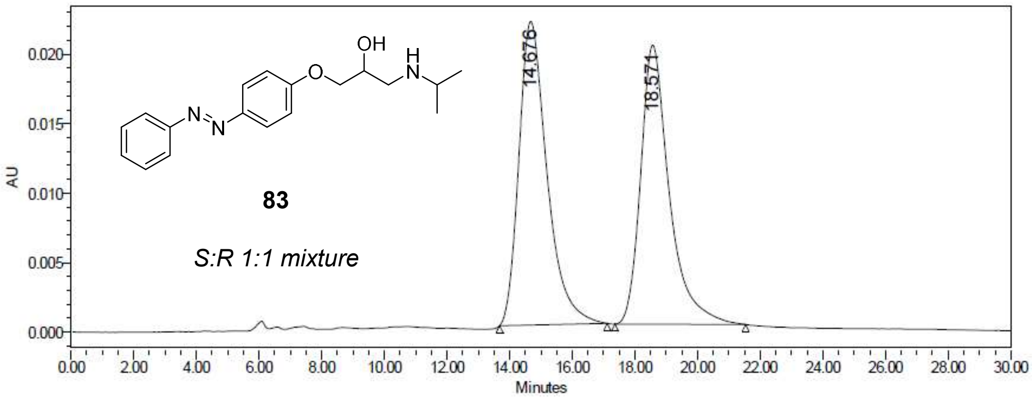

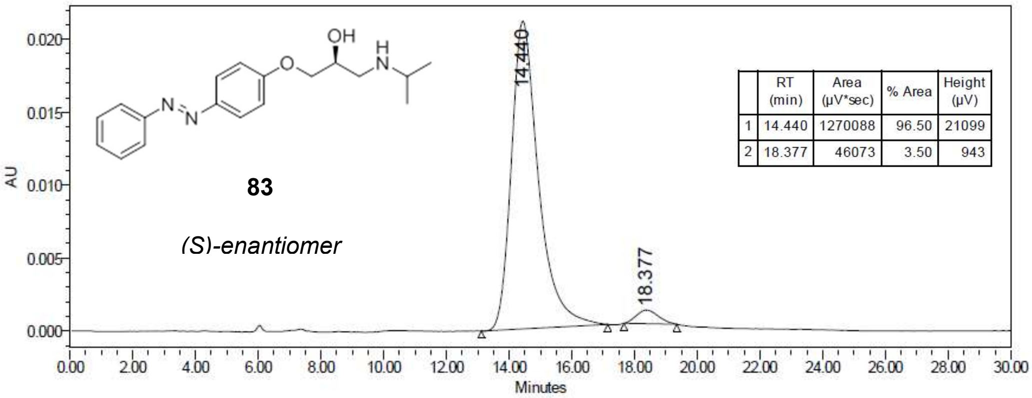

| | RT (min) | Area (µV*sec) | % Area | Height (µV) |
|---|---|---|---|---|
| 1 | 14.440 | 1270088 | 96.50 | 21099 |
| 2 | 18.377 | 46073 | 3.50 | 943 |

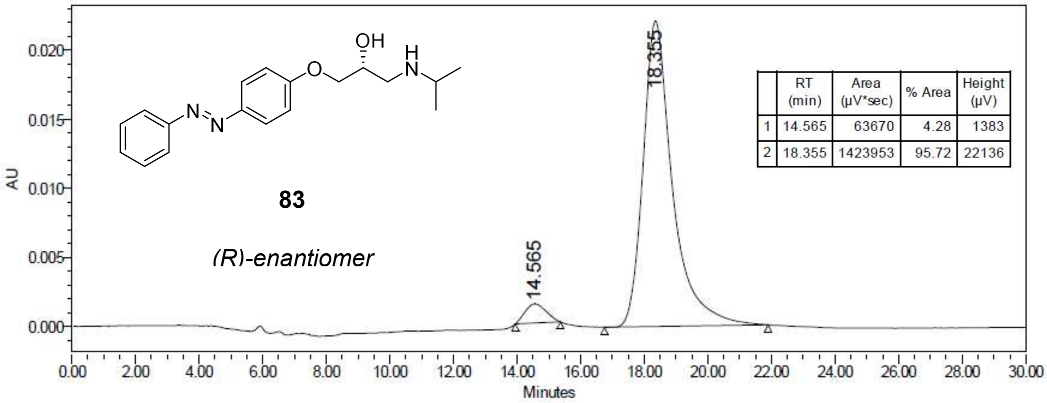

| | RT (min) | Area (µV*sec) | % Area | Height (µV) |
|---|---|---|---|---|
| 1 | 14.565 | 63670 | 4.28 | 1383 |
| 2 | 18.355 | 1423953 | 95.72 | 22136 |

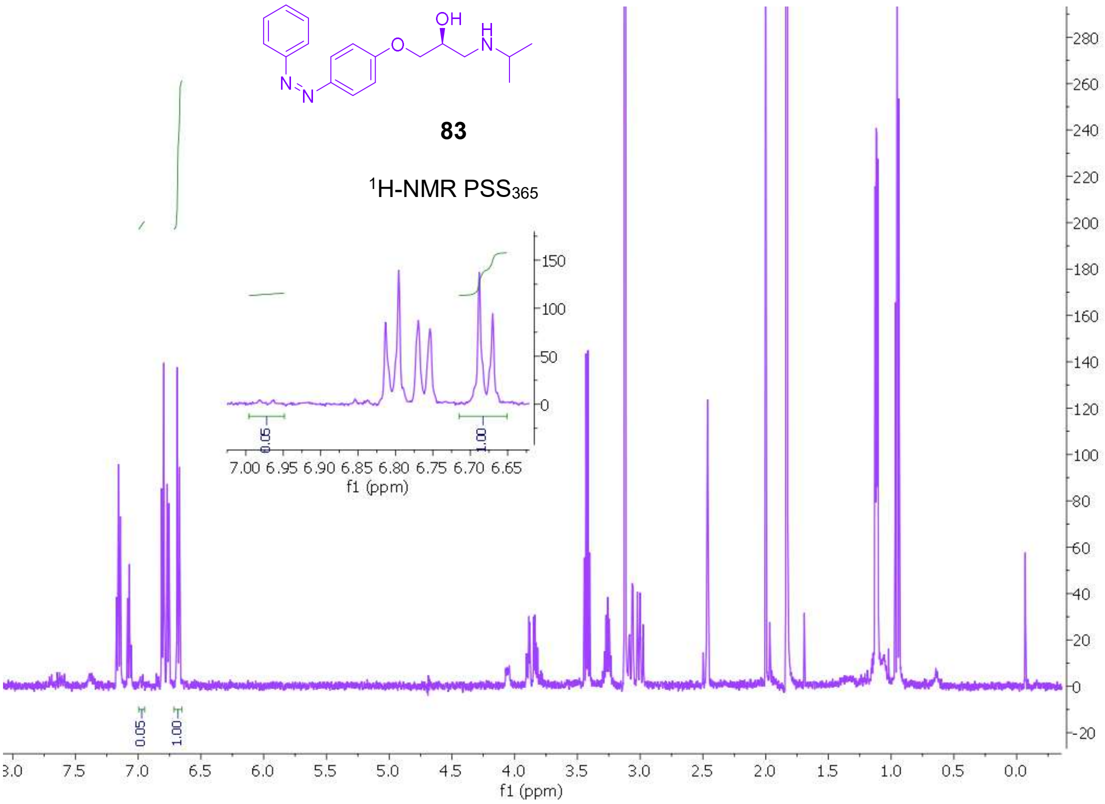
OH
H
N
O
N
N
83
1H-NMR PSS365
f1 (ppm)

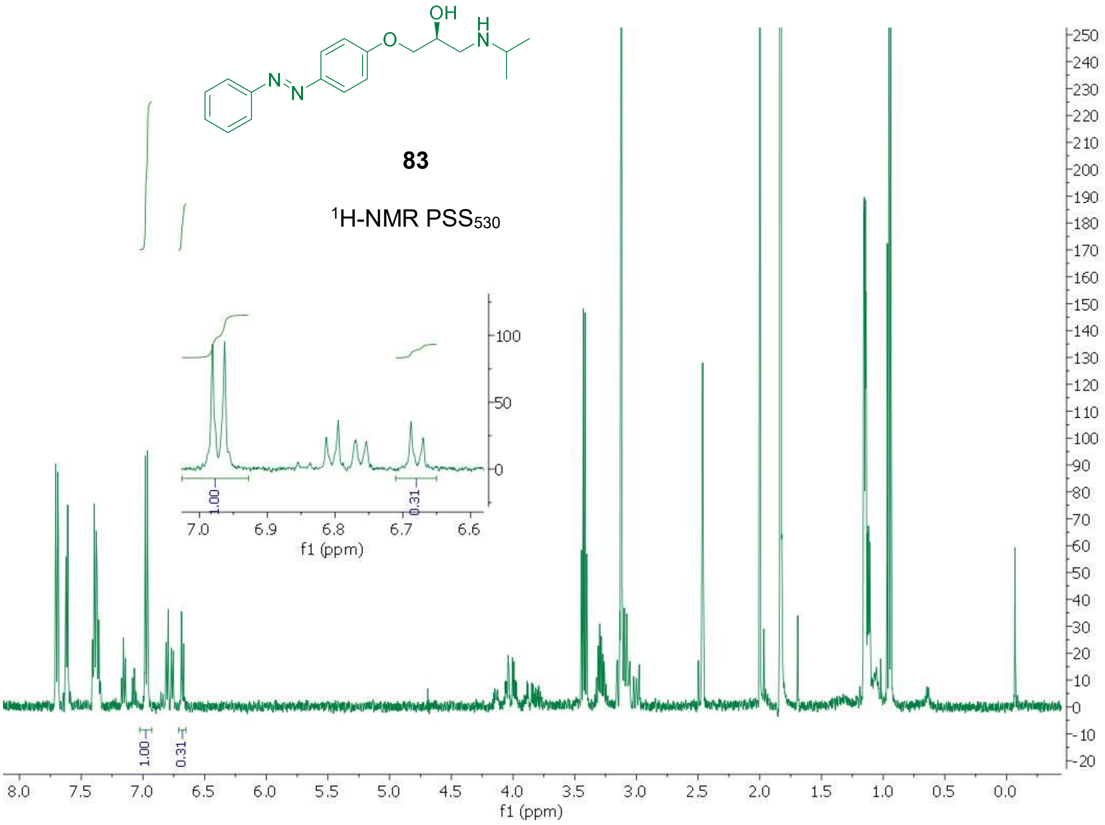
OH
H
N
O
N
N
83
1H-NMR PSS530
f1 (ppm)

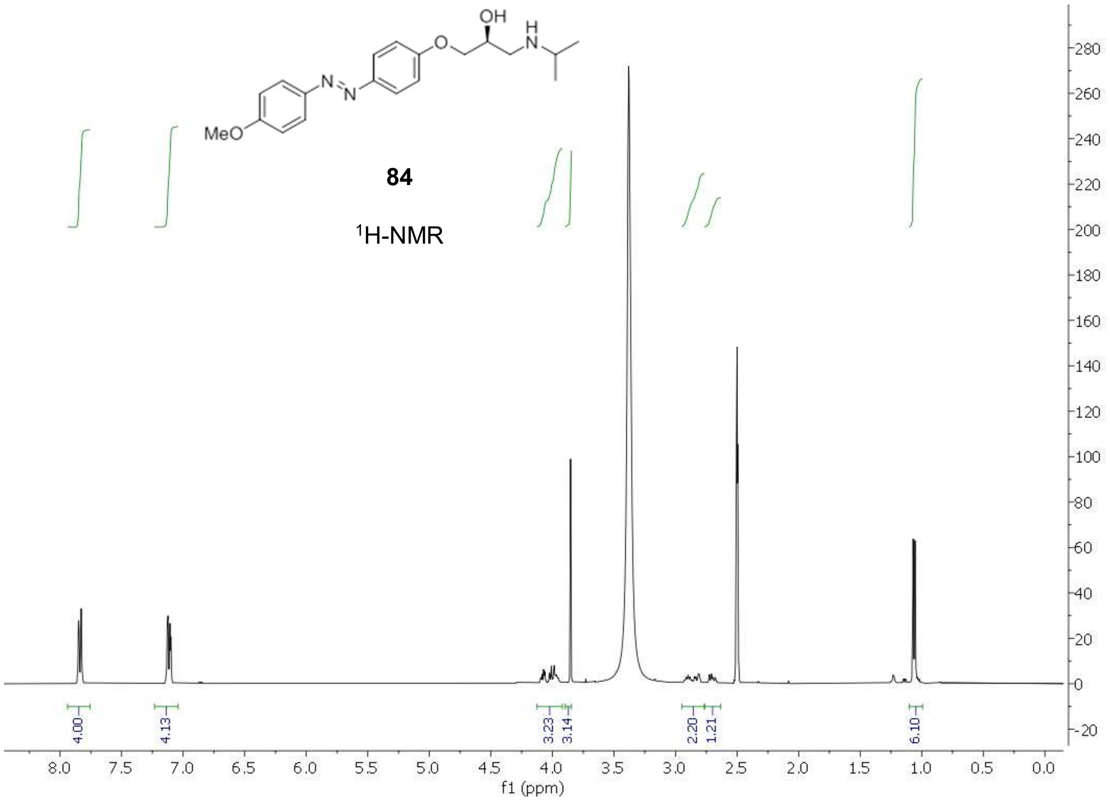

MeO
OH
H
N
84
1H-NMR
4.00
4.13
3.23
3.14
2.20
1.21
6.10
f1 (ppm)

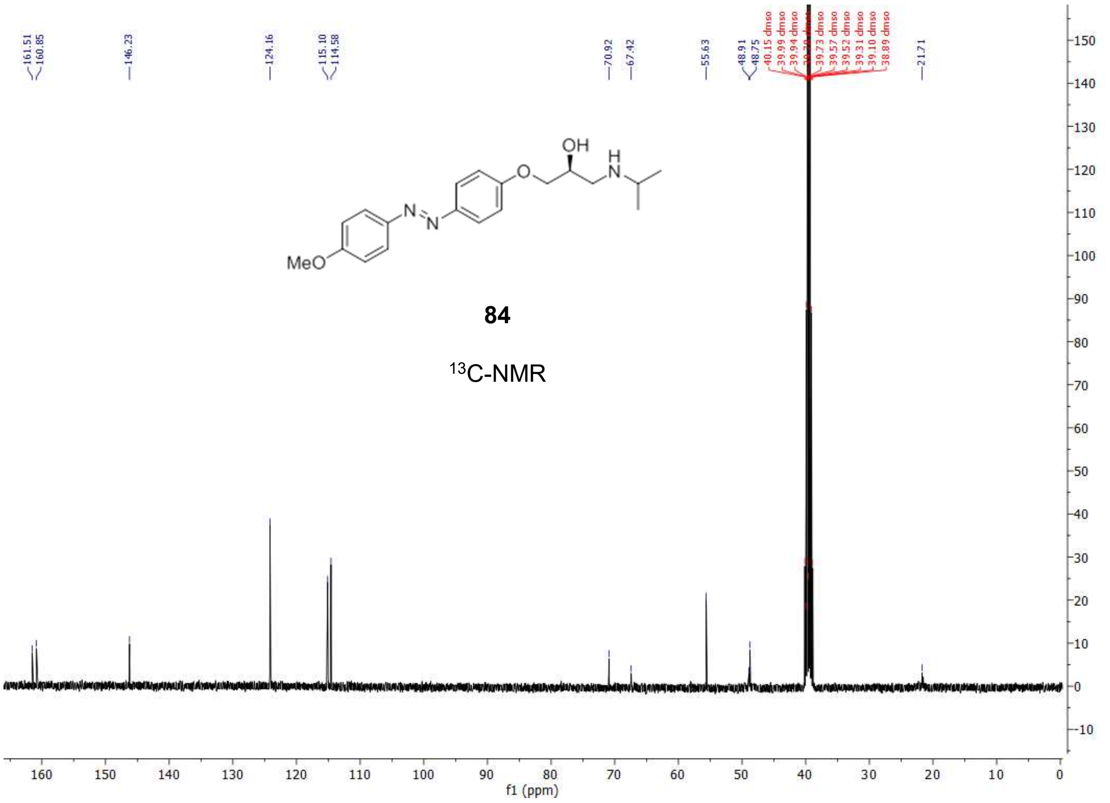

161.51
160.85
146.23
124.16
115.10
114.58
70.92
67.42
55.63
48.91
48.75
21.71
MeO
OH
H
N
84
13C-NMR
f1 (ppm)

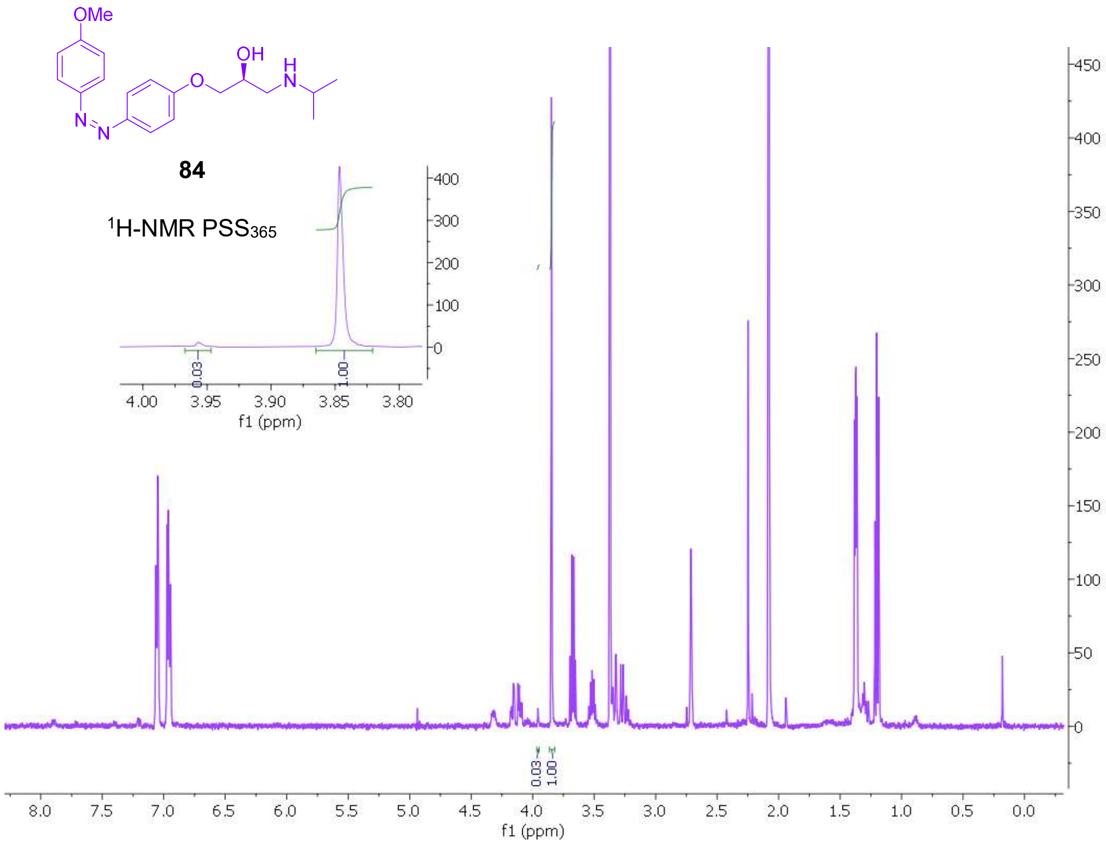
OMe
OH
H
N
N
N
O
84
1H-NMR PSS365
0.03
1.00
f1 (ppm)
f1 (ppm)

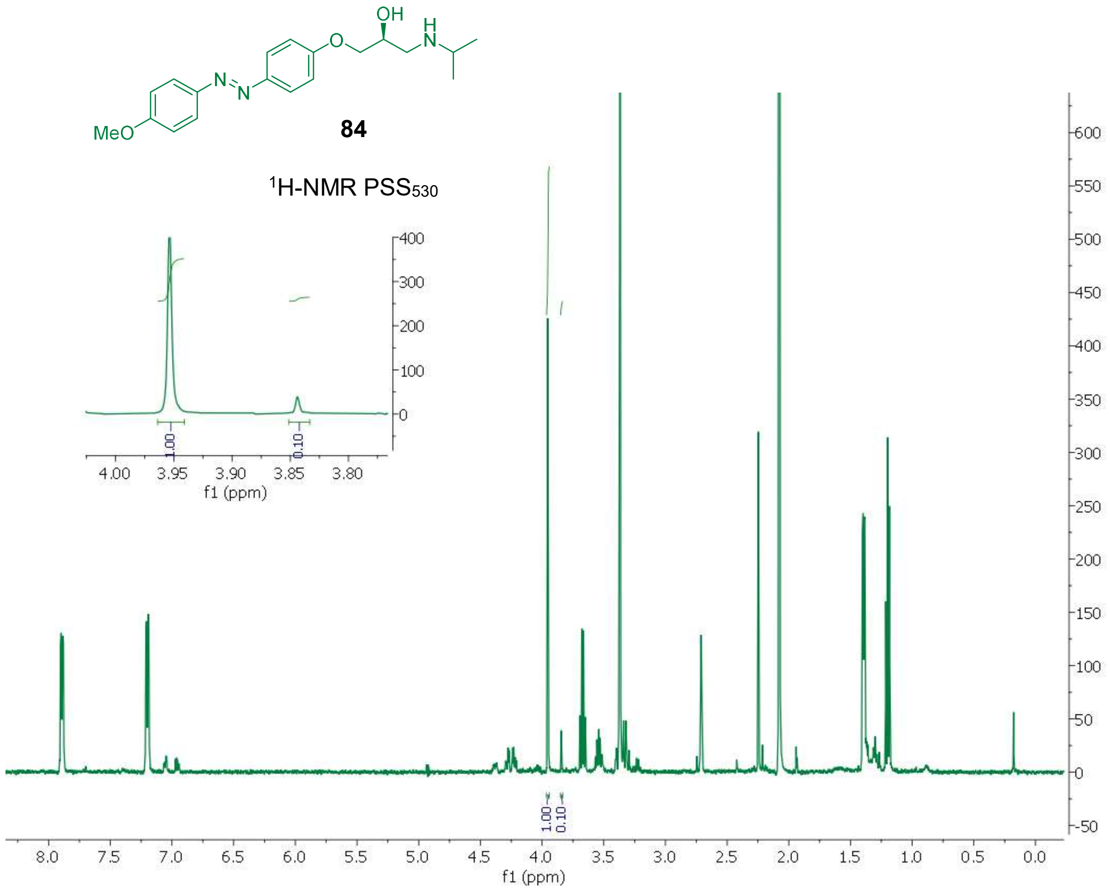
OH
H
N
O
N
N
MeO
84
1H-NMR PSS530
1.00
0.10
f1 (ppm)
f1 (ppm)
